# Designing and comparing cleaning pipelines for TMS-EEG data: a theoretical overview and practical example

**DOI:** 10.1101/2021.11.18.469167

**Authors:** Nigel C. Rogasch, Mana Biabani, Tuomas P. Mutanen

## Abstract

Combining transcranial magnetic stimulation (TMS) with electroencephalography (EEG) is growing in popularity as a method for probing the reactivity and connectivity of neural circuits in basic and clinical research. However, using EEG to measure the neural responses to TMS is challenging due to the unique artifacts introduced by combining the two techniques. In this paper, we overview the artifacts present in TMS-EEG data and the offline cleaning methods used to suppress these unwanted signals. We then describe how open science practices, including the development of open-source toolboxes designed for TMS-EEG analysis (e.g., TESA - the TMS-EEG signal analyser), have improved the availability and reproducibility of TMS-EEG cleaning methods. We provide theoretical and practical considerations for designing TMS-EEG cleaning pipelines and then give an example of how to compare different pipelines using TESA. We show that changing even a single step in a pipeline designed to suppress decay artifacts results in TMS-evoked potentials (TEPs) with small differences in amplitude and spatial topography. The variability in TEPs resulting from the choice of cleaning pipeline has important implications for comparing TMS-EEG findings between research groups which use different online and offline approaches. Finally, we discuss the challenges of validating cleaning pipelines and recommend that researchers compare outcomes from TMS-EEG experiments using multiple pipelines to ensure findings are not related to the choice of cleaning methods. We conclude that the continued improvement, availability, and validation of cleaning pipelines is essential to ensure TMS-EEG reaches its full potential as a method for studying human neurophysiology.

**Highlights:**

- Concurrent TMS-EEG is challenging due to artifacts in the recorded signals.
- We overview offline methods for cleaning TEPs and provide tips on pipeline design.
- We use TESA to compare pipelines and show changing a single step alters TEPs.
- We discuss the challenges in validating pipelines for TMS-EEG analysis.
- We suggest using multiple pipelines to minimise the impact of method choice on TEPs.

## 1. Introduction

Transcranial magnetic stimulation (TMS) is a non-invasive form of brain stimulation capable of depolarising and activating cortical neurons across the cortex using electromagnetic induction (Barker et al., 1985). Over the last several decades, various research groups have attempted to characterise the brain’s response to TMS by recording electroencephalography (EEG) during stimulation (Farzan et al., 2016; Ilmoniemi et al., 1997; Massimini et al., 2005; Rogasch and Fitzgerald, 2013). EEG uses electrodes placed on the scalp to measure the electric potential differences resulting from neural activity with high temporal resolution, thereby capturing the cascade of neural events occurring across the brain following TMS. The combination of TMS-EEG offers great promise for understanding how different cortical regions respond to TMS, how activity spreads across the brain following stimulation, and how different brain states and pathological conditions impact TMS-evoked brain activity (Tremblay et al., 2019). However, combining the two techniques is accompanied by several challenges. The TMS pulse causes a range of recording and physiological artifacts in EEG recordings which mask the neural activity evoked transcranially by the TMS pulse (Ilmoniemi and Kicić, 2010; Rogasch et al., 2014). While some of these artifacts can be avoided or minimised during data collection, others cannot. Furthermore, the presence of certain artifacts can interact with standard offline EEG-processing steps, resulting in additional analysis-related artifacts (Rogasch et al., 2017). Together, these factors have led to the development of offline artifact suppression methods and cleaning pipelines designed specifically for TMS-EEG data (Ilmoniemi et al., 2015).

The advent of bespoke cleaning approaches for TMS-EEG has led to its own series of challenges. Code for these novel approaches is often developed in-house, making it difficult for other research groups to replicate analysis pipelines or gain access to state-of-the-art cleaning methods. To meet this need, a number of open-source toolboxes have emerged which provide easy access to semi-standardised methods (Atluri et al., 2016; Rogasch et al., 2017). However, the choice of which methods to use and in which order leads to a combinatorial explosion of possible pipelines. A growing number of studies have shown that these choices matter, with different analysis pipelines resulting in discrepancies between the amplitude and spatial distribution of cleaned TMS-evoked potentials (TEPs) even when using the same data (Bertazzoli et al., 2021). Importantly, there is still no consensus on which pipeline to use, mainly due to the lack of ground truth for benchmarking pipelines against. Together, these challenges pose a daunting task for TMS-EEG researchers, who are faced with an ever-growing series of decisions which can ultimately impact the conclusions drawn from their data.

The aim of this paper is to provide an overview of the challenges associated with suppressing artifacts in TMS-EEG recordings, while providing practical recommendations on designing and implementing TMS-EEG analysis pipelines. We first introduce the challenges associated with recording neural activity using EEG following a TMS pulse due to artifacts, and some of the offline approaches designed to reduce the influence of artifacts in TMS-EEG data. We then overview the motivation and function of the TESA toolbox, an open-source library of MATLAB functions designed for cleaning TMS-EEG data which is implemented as a plugin in the EEGLAB toolbox. For comparison, we also contrast TESA against other open-source toolboxes available within the field. Next, we provide some general principles to consider when designing cleaning pipelines for TMS-EEG data. We then provide a practical example of how to use TESA to implement and test different cleaning pipelines by comparing different methods for removing the decay artifact following TMS. We show that even varying a single step in a TMS-EEG cleaning pipeline can result in different outcomes. Finally, we overview the challenges and future directions for validating TMS-EEG cleaning pipelines, before providing recommendations on how to minimise the impact of method choice on TMS-EEG findings.

## 2. Concurrent TMS-EEG: the challenge of artifacts

Recording EEG activity during TMS is challenging on several fronts. First, the TMS pulse can interact with the EEG recording equipment to induce several different types of artifacts. In this paper, we refer to artifacts as any EEG signal which is not generated as a direct result of transcranially stimulating the targeted cortical region with the TMS pulse. For our purposes, the signal of interest includes activity coming from stimulation of the target region and any transsynaptic activation of regions distant from the targeted region. The large time-varying magnetic field generated by a TMS pulse interacts with the skin-electrode-amplifier circuit to generate a large pulse artifact which is several magnitudes larger than neural activity measured with EEG (fig. 1A). Older EEG amplifiers were often saturated by these artifacts and only returned to useable recordings ranges after several seconds, making concurrent TMS-EEG impossible (Cracco et al., 1989; Izumi et al., 1997). More modern amplifiers (in the last two decades) have sufficient operating ranges to adequately capture the pulse artifact without saturation, recovering within 1-2 ms following the pulse (Bonato et al., 2006; Daskalakis et al., 2008; Rogasch et al., 2013; Veniero et al., 2009). Of note, care must be taken when setting the recording parameters of these amplifiers as wrong choices, e.g., in the cut-off frequency of the anti-aliasing filtering may lead to secondary artifacts, such ringing around the TMS-pulse artifact. Furthermore, additional transient artifacts with variable time lengths can occur in some amplifiers (fig. 1A). Other custom-designed amplifiers use a sample-and-hold circuit to pin the amplifier recording around the TMS pulse, thereby avoiding the artifact altogether (Ilmoniemi et al., 1997; Virtanen et al., 1999). The TMS pulse can also lead to additional charges or eddy currents which are stored at the skin/electrode interface, or possibly within the electrode lead wires and amplifier and result in ‘decay’ artifacts which broadly follow a power law distribution and can last from 10-100 ms following the TMS pulse (Freche et al., 2018; Julkunen et al., 2008; Sekiguchi et al., 2011) (fig. 1B). Contact between the TMS coil and electrodes can also introduce additional line noise (also known as radiofrequency noise) and movement artifacts (Ruddy et al., 2018) to these electrodes, and the recharging of the TMS capacitors following the pulse can also lead to a recharge artifact, the timing of which can vary in time and amplitude depending on the TMS intensity (Rogasch et al., 2013; Veniero et al., 2009). Further complicating matters, the presence and magnitude of these artifacts can depend on the hardware used to collect the data. While features of certain artifacts appear consistent between some hardware systems, there are published examples of changes in artifact profiles between different TMS devices (Rogasch et al., 2013), TMS coils (Fernandez et al., 2021), and EEG electrodes (Mancuso et al., 2021), as well as anecdotal evidence within the TMS-EEG community for differences between EEG amplifiers. The vast number of different hardware combinations used by different laboratories poses a genuine challenge for comparing data between groups, which may suffer from different, and in some cases unique, artifact profiles.

**Figure 1:**
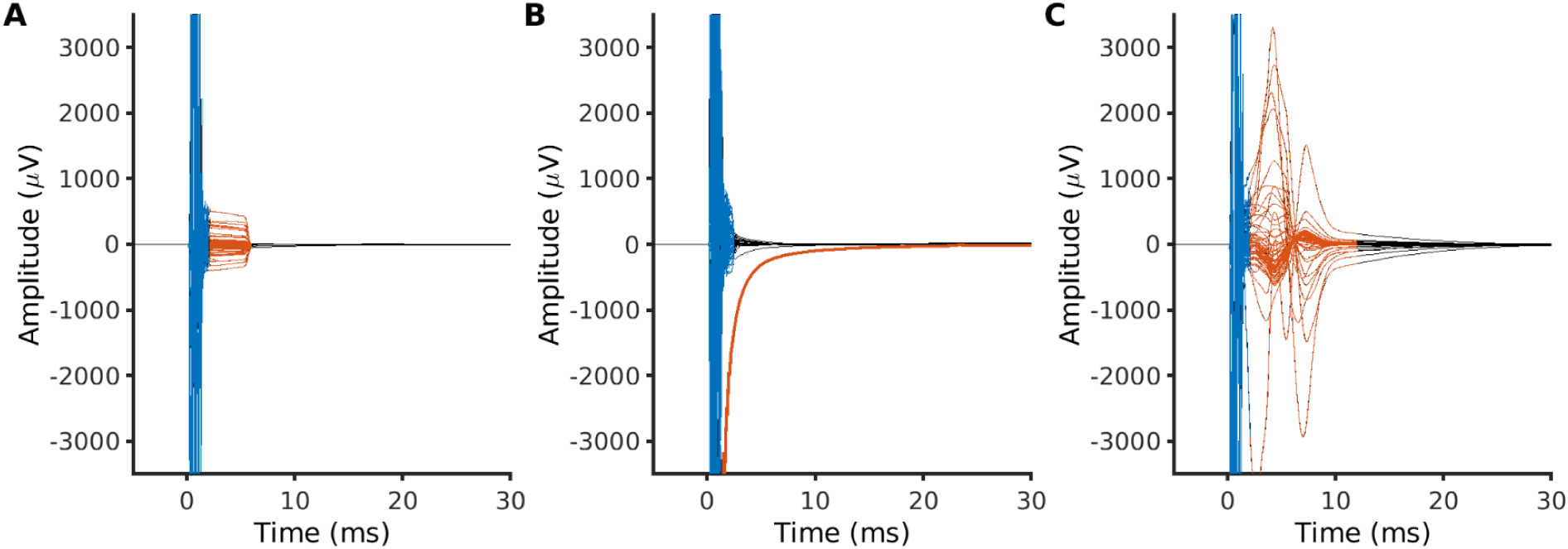
Common artifacts in TMS-EEG recordings. A) An example of the TMS pulse artifact (blue) lasting from 0-2 ms followed by an additional transient artifact likely related to the amplifier (red) lasting until ∼6 ms. B) An example of an electrode with a large decay artifact (red). C) An example of a TMS-evoked muscle artifact (red) with peaks at ∼4 ms and ∼8 ms.

The second major cause of contamination in TMS-EEG recordings are EEG signals resulting from the interaction between the TMS pulse and physiological systems. The TMS pulse can depolarise face and scalp muscles either directly or indirectly through activation of motor neurons innervating the muscles (Korhonen et al., 2011; Mutanen et al., 2013; Rogasch et al., 2013). The resulting compound muscle action potential is similar in shape and size to M-waves recorded from peripheral muscles following transcutaneous nerve stimulation or motor-evoked potentials (MEPs) following TMS, and typically shows bipolar peaks at ∼4-5 and 8-10 ms following the pulse which last for 10s of ms (note the exact timing depends on the EEG sampling rate and filter settings) (fig. 1C) (Mutanen et al., 2013; Rogasch et al., 2013). TMS-evoked muscle activity becomes more prominent over lateral stimulation sites and can reach peak-to-peak amplitudes an order of magnitude larger than neural signals in the EEG (>1-2 mV) (Mutanen et al., 2013; Rogasch et al., 2013). Movement associated with TMS-evoked scalp muscle contraction can cause additional decay artifacts which persist for longer (>50 ms) and are often hard to distinguish from decay artifacts associated with charges stored within the skin-electrode-amplifier circuit (Rogasch et al., 2014). The often-strong sensations resulting from TMS-evoked muscle activity or from stimulation of other cranial nerves running across the scalp, may also result in somatosensory-evoked potentials (Conde et al., 2019; Gordon et al., 2021). There is some debate as to whether these potentials are directly visible in the scalp EEG (Paus et al., 2001; Rocchi et al., 2021), however other work has shown an increase in potentials around 100-200 ms with increasing electrical stimulation to the scalp (Gordon et al., 2021). The discomfort caused by TMS-evoked muscle activity can also result in jaw or facial tension, which causes high levels of ongoing muscle activity in the EEG signal, particularly in lateral electrodes overlying muscles. Furthermore, TMS pulses over frontal region can trigger a blink response in some individuals, which is often tightly linked to the stimulus, occurring 80-150 ms following the TMS pulse and causing deflections >20-30 µV in amplitude in frontal electrodes (Bruckmann et al., 2012; Rogasch et al., 2014, 2013). Finally, TMS results in a loud clicking sound, which evokes a prominent auditory potential with peaks at ∼100 and 200 ms (Nikouline et al., 1999). Importantly, sensory-evoked potentials can persist in TMS-EEG recordings even when online masking methods are used, like playing noise through earphones to mask the TMS click (Biabani et al., 2021, 2019; Conde et al., 2019; Fernandez et al., 2021; Gordon et al., 2018; Herring et al., 2015). Of note, whether sensory-evoked potentials like those resulting from the TMS coil click are considered artifacts is largely dependent on the goal of the study. If one is simply interested in quantifying the brain’s response to TMS, then sensory-evoked responses are a legitimate neural response to the stimulation which will be present in all TMS experiments that do not minimise sensory input (i.e., the vast majority of the TMS literature). However, if one is specifically interested in the neural response from the cortical circuits stimulated transcranially by the TMS pulse, then sensory-evoked potentials are a problem which potentially mask or interact with this TMS-evoked activity. Together, these different sources of artifacts can account for >70% of the variance in the data with certain experimental arrangements (after excluding the TMS pulse artifact) (Rogasch et al., 2014), substantially masking the patterns of neural activity directly resulting from the transcranial effects of stimulation on cortical circuits. For a detailed discussion on artifacts in TMS-EEG, please see other reviews (Ilmoniemi et al., 2015; Ilmoniemi and Kicić, 2010; Rogasch et al., 2017) .

## 3. Minimising artifacts in TMS-EEG data: online and offline approaches

Given the substantial contribution artifacts can make to TMS-EEG signals, a broad spectrum of approaches have emerged across the TMS-EEG community for minimising artifacts. At one end of the spectrum, online approaches attempt to avoid all artifacts during data collection. At the other end of the spectrum, offline approaches do not try to avoid artifacts during recording and instead attempt to remove or suppress artifacts using different data cleaning approaches. In reality, most groups use varying combinations of online and offline methods for minimising artifacts in TMS-EEG data. These variations likely reflect genuine differences in the recording setups and research questions across different groups. However, the large variability in online and offline approaches between studies also likely underlies the large variability in TEP characteristics reported throughout the literature (e.g., amplitude, shape, spatial distribution), as noted by several recent commentaries on the state of the TMS-EEG field (Belardinelli et al., 2019; Siebner et al., 2019). Perhaps the greatest challenge facing the field is the development of unified testing and analysis procedures that allow more direct comparisons of TEPs across research groups.

The online approach can include a variety of methods employed during data collection, such as: using equipment that is robust to TMS pulse and decay artifacts (e.g., TMS coils, EEG amplifiers, EEG electrodes); careful preparation of electrode-skin interfaces to minimise impedance and avoid decay artifacts (this can include rotating electrode leads to maximise common-mode rejection (Sekiguchi et al., 2011), or even micropuncture of the skin (Julkunen et al., 2008)); delaying the capacitor recharge of the TMS device to time periods outside the analysis window (Rogasch et al., 2013)(note that toolboxes like MAGIC allow external control of TMS device stimulation settings including capacitor recharge timing (Habibollahi Saatlou et al., 2018)); only stimulating cortical regions close to midline to avoid TMS-evoked muscle activity, blinks and other strong sensations (Ferrarelli et al., 2012; Massimini et al., 2005; Rosanova et al., 2009); and using noise-masking techniques which match the spectral properties of the TMS clicking noise as closely as possible (Massimini et al., 2005; Russo et al., 2021). Importantly, this approach also involves checking the TEP signals online and adjusting the coil position/orientation and stimulation intensity to 1) ensure a large TEP response within the first 50 ms which is thought to represent activity from the stimulated site; and 2) ensure TMS-evoked muscle and other artifacts are reduced (reviewed in preprint: Casarotto et al., 2021). The major advantage of this approach is that it ensures excellent signal-to-noise ratios and greatly simplifies offline cleaning pipelines. However, this approach is only suited to certain experimental designs which involve stimulation of midline cortical sites and allow repositioning of the TMS coil within a certain radius. Furthermore, until recently there were few open-source or commercial tools available that allowed checking TEPs online, although several toolboxes have been recently developed (e.g., rt-TEP toolbox (Casarotto et al., 2021); BEST toolbox (Hassan et al., 2021)).

The offline approach has led to the development or modification of a range of EEG analysis methods specifically for targeting artifacts common in TMS-EEG signals. For example, blind source separation algorithms, including independent component analysis (ICA; approaches include INFOMAX, FastICA, and enhanced deflation method) (Hamidi et al., 2010; Hernandez-Pavon et al., 2012; Korhonen et al., 2011; Metsomaa et al., 2014; Rogasch et al., 2014), and principal component analysis (PCA; approaches include data compression and suppression) (Hernandez-Pavon et al., 2012; ter Braack et al., 2013) have been modified for identifying and suppressing the TMS-evoked muscle artifact, as have source-based spatial filtering methods (signal-space projection [SSP], and SSP with source-informed reconstruction [SSP-SIR]) (Mäki and Ilmoniemi, 2011; Mutanen et al., 2016). Various methods have been developed for suppressing decay artifacts including the ICA variants mentioned above, the source-estimate-utilising noise-discarding (SOUND) algorithm which uses iterative Wiener estimation and EEG forward modelling to detect and suppress noise from individual electrodes (Mutanen et al., 2018), and methods which fit different models to the decay signal (e.g., linear functions, exponential functions, power-law functions, or physical models of the skin-gel-electrode interface) and then subtract the model estimates from the data (Casula et al., 2017; Freche et al., 2018; Van Der Werf and Paus, 2006; Zanon et al., 2010). ICA is also typically used to suppress other common artifacts like eye blinks/movement and ongoing muscle activity (Rogasch et al., 2014), whereas ICA, SSP-SIR, and template subtraction (e.g., subtracting a control condition) have been trialled to separate TMS-evoked sensory activity from activity resulting from the transcranial stimulation of cortical circuits (Bender et al., 2005; Biabani et al., 2019; Rogasch et al., 2014). For a detailed description of different cleaning methods, the reader is directed to other reviews (Ilmoniemi et al., 2015; Ilmoniemi and Kicić, 2010; Rogasch et al., 2017).

The major advantage of the offline approach is that the choice of stimulation location is not limited to midline areas away from scalp/facial muscles. However, there are a number of challenges associated with suppressing artifacts offline. First, many of the novel analysis methods for cleaning TMS-EEG data are developed in-house, meaning it is difficult for most users to reproduce the code required to implement these analyses, thus leading to a reproducibility issue. Second, the large number of different analysis approaches leads to an explosion of possible cleaning pipeline combinations. Of particular note, validating these different analysis methods and pipelines (i.e., determining whether they accurately suppress artifacts while retaining the signal of interest) is extremely difficult as the ‘ground-truth’ signal of interest (i.e., the TMS-evoked neural response) is unknown. Third, combining multiple preprocessing steps into a complicated pipeline adds the possibility for undesired interaction effects on the cleaned signal that are difficult to control. Therefore, researchers are faced with a series of difficult decisions of which analysis pipeline to choose and are ultimately unsure whether the resulting ‘cleaned’ TEPs are an accurate reflection of the neural signal of interest. We will discuss each of these challenges in more detail in the following sections.

## 4. The TESA toolbox

To overcome the reproducibility challenges associated with offline TMS-EEG analysis, several open-source TMS-EEG analysis toolboxes have become available over the last 5 years. One of these led by our team is TESA (TMS-EEG signal analyser) (Rogasch et al., 2017). TESA was designed with two main goals: 1) to provide a standardised library of offline analysis methods used in TMS-EEG research, and 2) to make TMS-EEG analysis accessible to researchers without a scripting/coding background. To achieve these outcomes, TESA was built as a plugin (also called an extension) to EEGLAB (Delorme and Makeig, 2004), the popular open-source EEG analysis toolbox run on the MATLAB platform. Building TESA within EEGLAB offers several key advantages. First, EEGLAB already has a large library of functions for EEG analysis which are available for use with TESA functions and is familiar to many researchers already undertaking EEG research. Second, EEGLAB has a modular framework which allows for flexibility in designing and implementing analysis pipelines. Finally, EEGLAB has a well-established graphical user interface (GUI) which makes the practicalities of running analysis much more intuitive for users not familiar with coding. For those less proficient in programming, the EEGLAB GUI allows an easy transformation of the history of the past preprocessing steps into an executable MATLAB script for later replication of the analysis. TESA is accompanied by a free online user manual, which details both the theory underlying the analysis functions in TESA, and detailed instructions on how to use the toolbox (https://nigelrogasch.gitbook.io/tesa-user-manual/). The TESA user manual forms an integral part of educating researchers on the progress and pitfalls of TMS-EEG analysis.

In recognition that there is no clear consensus on the best way to analyse TMS-EEG data, TESA does not advocate for any one TMS-EEG processing pipeline. Indeed, one of the central motivations for TESA is to provide access to as many analysis functions as possible, thereby allowing comparison of how different approaches impact the outcome of TMS-EEG analysis. TESA is continually evolving as new methods become available. Recent additions include source-based artifact cleaning methods such as SSP-SIR and SOUND (Mutanen et al., 2020), as well as fit and remove functions included in the most recent release accompanying this article (Freche et al., 2018). Importantly, TESA is hosted on github, a code repository which allows users to both download and contribute to the toolbox (https://github.com/nigelrogasch/TESA/releases). It is hoped that TESA will continue to grow and evolve with the help of the TMS-EEG community.

In addition to TESA, a range of other open-source TMS-EEG toolboxes are also available with differing principles and functionality. TMSEEG is a dedicated TMS-EEG analysis toolbox also designed within the EEGLAB framework but has a more standardised workflow and a dedicated GUI (Atluri et al., 2016). TMSEEG offers some additional features such as channel removal and interpolation at the single trial level, which can be useful for minimising data loss, but on the other hand lacks some artifact-rejection techniques, complimentary to ICA. FieldTrip, another popular open-source EEG toolbox in MATLAB with similar broad functionality to EEGLAB but no GUI (Oostenveld et al., 2011), has several functions dedicated to TMS-EEG analysis which integrate within the modular design of the toolbox (https://www.fieldtriptoolbox.org/tutorial/tms-eeg/). ARTIST is a stand-alone TMS-EEG analysis toolbox implemented in MATLAB (Wu et al., 2018). Unlike other toolboxes, ARTIST is designed as a fully automated TMS-EEG analysis pipeline which uses a combination of heuristic rules and machine learning approaches to automate manual steps in the analysis (e.g., selecting trials/channels/ICA components for removal). To facilitate automation, the pipeline of ARTIST is fixed, and restricted to data sets with similar electrode layouts due to the classifiers used to select ICA components. In addition to specific toolboxes, several novel TMS-EEG cleaning functions and pipelines have been developed and the code made publicly available, which is another excellent approach for ensuring reproducibility (Cline et al., 2021; Freche et al., 2018). By developing and sharing toolboxes and code, the open-source approach has largely overcome the analysis reproducibility issue facing TMS-EEG research, and there are now a number of excellent options available to TMS-EEG researchers for accessing state-of-the-art cleaning approaches. However, the large number of open-source options has not yet solved the problem of variability in preprocessing pipelines across research groups, which still hinders comparisons of TMS-EEG results within the community.

## 5. Considerations for designing and implementing TMS-EEG cleaning pipelines

Open-source toolboxes have provided access to a range of analysis functions designed specifically for TMS-EEG data cleaning. The next consideration is how these cleaning steps fit within an overall TMS-EEG analysis pipeline. The unique artifact profile of TMS-EEG data means that the order of analysis steps typically used for standard event-related EEG paradigms are often not appropriate. Importantly, certain cleaning steps can introduce additional analysis-related artifacts if used inappropriately with TMS-EEG data (Rogasch et al., 2017). It is also possible that performing certain preprocessing steps in a suboptimal order can cause complicated interaction effects on the final outcome. Furthermore, the assumptions underlying certain cleaning steps used in EEG data may not hold as strongly for TMS-EEG data. In this section, we outline some of the key considerations when designing and implementing TMS-EEG cleaning pipelines, as well as providing some practical tips to help identify issues when they arise.

### 5.1. Minimise artifacts during data collection

Given the challenges in removing artifacts offline, all efforts should be made to avoid artifacts during TMS-EEG data collection. While this recommendation does not directly refer to the design of cleaning pipelines per se, raw data which contains less artifacts is considerably easier to clean and requires less assumptions, thereby greatly simplifying data analysis. At a minimum, it is recommended to ensure electrode-skin preparation is as good as possible, with low impedance values maintained throughout the experiment (preferably <5 kΩ) to minimise decay artifacts. For EEG caps which allow rotation of electrodes, adjusting the orientation of the electrode wires may also help optimise cancellation of decay artifacts. Participants should be encouraged to keep facial muscles as relaxed as possible during the experiment to avoid ongoing muscle artifacts. Some form of feedback from the experimenter when muscle activity is present (e.g., a tap on the shoulder or visual cue) may aid in minimising muscle activity. The researchers should also ensure a maximally relaxed position of the participant and avoid pressing the stimulating coil against the head, to minimize any neck and head muscle tension. Furthermore, TMS capacitor recharge should be delayed to a time point not within the time range of interest (usually within 500 ms of the TMS pulse) when possible. Wherever the experimental design allows, midline stimulation sites which do not evoke muscle activity or blinks following TMS should be preferred. Ideally the signals should be checked online to ensure TMS-evoked muscle and blink artifacts are not present in the data, and the coil position adjusted until these artifacts are minimised (Mutanen et al., 2013). Finally, every step should be taken to minimise auditory input following the TMS pulse to avoid auditory-evoked potentials (Rocchi et al., 2021; Russo et al., 2021), including using a barrier between the coil and electrodes to minimise vibration (Ilmoniemi et al., 2015). Taking these steps might add additional time to the experiment but can substantially simplify the cleaning process, thereby increasing confidence in the final TEPs.

### 5.2. Design a pipeline appropriate for the artifacts present in the data

It is tempting to think that a goal of TMS-EEG pipeline development should be to design a single validated cleaning pipeline that is used ubiquitously in TMS-EEG research. However, the large variation in artifact profiles present in TMS-EEG data means that it is highly unlikely a single pipeline will be appropriate for all cases. For example, a study design that allows coil positioning away from scalp muscles can result in a relatively ‘clean’ signal that would require a minimal cleaning pipeline that does not require some of the more aggressive cleaning approaches. In contrast, a study design that requires stimulation of lateral areas like the dorsolateral prefrontal cortex will inevitably contain TMS-evoked muscle activity and will therefore require a specialised pipeline for adequately suppressing this artifact source to uncover the underlying neural activity. Moreover, different TMS-EEG setups capture different artifacts, naturally leading to particular preprocessing requirements. Given that signal separation methods like ICA and SSP cause substantial alterations to the data, these should only be used when absolutely necessary, and ideally avoided if possible (see recommendation 4). Therefore, it is important to carefully consider the artifact profile of the data when designing TMS-EEG cleaning pipelines, and only use ‘aggressive’ cleaning steps like signal separation methods if required.

### 5.3. Avoid using temporal filters over large steps in the data

Perhaps the most important consideration when designing TMS-EEG cleaning pipelines is minimising large voltage steps or sharp transitions in the data prior to the use of temporal filters. Large voltage steps are caused by a number of factors in TMS-EEG experiments. First, the TMS pulse causes a very high amplitude and high frequency artifact, which is short-lasting and typically removed from the data. Beyond the pulse artifact, the post stimulation signal can suffer from long lasting offsets from the pre stimulation signal due to decay and/or TMS-evoked muscle artifacts. Finally, the recharging of the TMS capacitors can also result in a high-frequency artifact. Temporal filters are commonly applied to EEG data to attenuate the contribution of low-frequency (high-pass filter) or high-frequency (low-pass filter) noise to the data (or both at the same time: band-pass filter), to prevent aliasing artifacts caused by downsampling (anti-aliasing filter), or to attenuate narrow-band noise such as line noise (band-stop or notch filters) (de Cheveigné and Nelken, 2019). Temporal filtering is also useful for optimising other processing steps like ICA, the convergence of which can suffer from low frequency drifts in the signal (Winkler et al., 2015). However, applying temporal filters over voltage steps or sharp transitions can introduce unwanted distortions to the data such as ringing or drift artifacts due to the interaction between the filter and these features in the signal (de Cheveigné and Nelken, 2019). Importantly, these spurious signals can last up to several 100 ms and resemble the shape of oscillations and event-related potentials (ERPs). As acausal, ‘zero-phase’ filters are often used in EEG analysis (i.e., the filter is passed forward and backward through the data to minimise phase offsets), filtering artifacts can be introduced both before and after the transition event.

Given that TMS introduces voltage steps and high-frequency events into the signal, it is imperative that these artifacts are removed or suppressed as much as possible prior to any temporal filtering (anti-aliasing prior to downsampling, band-pass or band-stop filters) to prevent the introduction of ringing artifacts. Removing the data impacted by the TMS pulse artfiact and interpolating the missing data using a cubic or autoregressive functions to smooth the transition between the pre and post signals can minimise ringing from antialiasing filters used prior to downsampling (Cline et al., 2021; Rogasch et al., 2017). If TMS-evoked muscle or decay artifacts are present in the data, ICA, SOUND, or fit-and-subtract methods provides some benefit prior to band-pass filtering by attenuating the large amplitude steps associated with these artifacts. Furthermore, the design of the filter (e.g., the steepness of the transition in the frequency domain governed by the order of the filter) may also require optimisation, although we are unaware of any systematic comparisons between filter designs for TMS-EEG data. A recent study has suggested a modified high-pass filter to avoid ringing/drift artifacts around the TMS pulse, which uses an autoregressive model to extend the pre and post artifact data forward and backward respectively, and then filters each section separately (Cline et al., 2021).

It is noteworthy that some anti-aliasing filtering is always needed in the EEG amplifier before digitising the data at the given sampling rate. Because many of the abrupt TMS-pulse-related artifacts cannot be avoided during the experiments, the sampling rate should be set sufficiently high to avoid ringing in the measured signals due to the amplifier filters. For instance, if the EEG amplifier does not have the sample-and-hold circuitry to block the TMS-pulse artifact, a sampling rate of 5000 Hz or higher (allowing a low-pass filter of at least 1000 Hz) ensures a short-lived disturbance during the magnetic stimulus. It is also a common approach to high-pass filter the TMS-EEG data to remove slow drifts in the data . If the data consist of several ocular artifacts, which last hundreds of milliseconds, high-pass filtering before the removal of these sporadic artifacts may lead to additional low-frequency ringing artifacts. Therefore, using a minimal high-pass filter (or no high-pass filter if using a DC-coupled amplifier) during data recording is also recommended.

Another important consideration resulting from the necessity to use certain steps prior to temporal filters is that some of these methods require the data to be epoched. Filtering over epoched data can introduce additional artifacts at the epoch boundaries caused by zero-padding the data. As such, it is important to ensure that epochs are of sufficient length to discard affected data without corrupting the time periods of interest (e.g., the times around the TMS pulse). The take home message is that the order in which analysis steps are performed matters for TMS-EEG data because of the unique artifact profile caused by combining the techniques. Careful consideration for how different analysis steps interact with one another is therefore necessary when designing TMS-EEG analysis pipelines.

### 5.4. Use signal separation approaches with caution

ICA is a signal separation method used nearly ubiquitously in TMS-EEG data cleaning pipelines to suppress artifact sources within the data. Indeed, applying ICA to experimental data results in components which have topographies and time courses consistent with neural and artifacts sources expected within TMS-EEG data (Rogasch et al., 2014). However, an important question which is often overlooked is how accurately these components reflect the underlying signals. An important assumption of ICA is that the underlying signals are statistically independent. Practically, this assumption requires that there is sufficient temporal variability between each independent source underlying the EEG signal. TMS causes highly time-locked activity from both neural and artifactual signals, substantially reducing temporal variability and thereby weakening the statistical independence between sources. Metsomaa and colleagues (Metsomaa et al., 2014) demonstrated this problem using simulated TMS-EEG data, showing that standard applications of FastICA failed to accurately decompose the spatial topographies of most of the underlying signals in the presence of typical TMS-evoked artifacts (e.g., TMS-evoked muscle and decay artifacts). Similar results were reported by Hernandez-Pavon, Metsomaa and colleagues (Hernandez-Pavon et al., 2012) when trying to recover the time course and topography of a simulated signal combined with real TMS-EEG data, particularly in the presence of large TMS-evoked muscle artifacts. Together, these studies suggest that while appearing plausible, the component topographies following ICA of TMS-EEG data may reflect inaccurate representations of the underlying signals. The practical implication is that removing components from the data, as is done when ICA is used to identify and remove artifacts, can result in distortion of the underlying neural signals due to inaccurate ICA decomposition, especially when large signals like TMS-evoked muscle activity are present in the data. Reducing the size of TMS-evoked muscle activity either using PCA to suppress the affected segments of the signal (Hernandez-Pavon et al., 2012), or by removing the main peaks of the muscle activity (e.g., the first 10-15 ms of data) (Rogasch et al., 2014) may partially help with this problem, although it remains unclear to what extent.

The issue of data distortion is not limited to ICA. Other signal separation methods like SSP also cause substantial alterations to the underlying signals. For example, Mäki and colleagues (Mäki and Ilmoniemi, 2011) showed that SSP could help identify and suppress TMS-evoked muscle activity by using the high-frequency (>100 Hz) part of the signal to identify topographies related to muscle activation, however this approach also severely attenuated the remaining EEG signal, especially in electrodes near the site of stimulation. Mutanen and colleagues (Mutanen et al., 2016) showed that using a source-informed reconstruction approach (SSP-SIR), which relies on accurate modelling of the lead-field covariance matrix, can help with the attenuation, although this approach does not fully compensate the overcorrection of neuronal signals when large-amplitude artifacts are suppressed. Biabani and colleagues (Biabani et al., 2019) showed the SSP-SIR approach could also help in separating sensory-evoked signals from TMS-evoked signals in the TEP and resulted in more plausible sensor and source patterns compared with linear regression and ICA. However, the remaining signal was severely attenuated in amplitude and it remains unclear whether the resulting TEP was an accurate representation of the strength of the underlying TMS-evoked neural activity. An important assumption of SSP is that the artifact of interest is spatially uncorrelated with the transcranially-evoked neural activity, an assumption which is not likely to hold for all experimental arrangements (e.g., if the site of stimulation overlaps with regions activated by the sensory input following TMS). Together, these caveats suggest that caution is required when interpreting TMS-EEG data following the use of signal separation methods, as these methods can substantially distort the neural signals of interest while suppressing artifact signals.

### 5.5. Concatenate data sets where appropriate

Often, TMS-EEG experiments involve the comparison of TEPs collected from different experimental conditions (e.g., comparing TEPs from stimulating the same site over different time points before and after an intervention; or comparing TEPs from stimulating different cortical sites). One approach that is sometimes used in these experiments is to concatenate the different conditions at the beginning of the analysis pipeline (usually after epoching) (Chung et al., 2017; Rogasch et al., 2015). The main rationale for concatenating data sets is that if signal separation steps like ICA are used on the data, component removal is applied equally across conditions, therefore preventing any unintended biases from inconsistently removing components if the conditions are analysed separately. Furthermore, including more data points can help meet the minimum data requirements for running ICA. However, concatenating data sets may not always be appropriate. For example, ICA decomposition may be sub-optimal if there are substantially different signal profiles (both brain and artifact) between conditions. This is often the case when stimulating different cortical sites, which substantially changes the artifact profile and may activate different neural populations. ICA should not be run over data sets where the electrode positions have changed (e.g., from reapplying the cap, or even re-gelling the electrodes mid experiment), as this can invalidate the assumption of spatial stationarity within the data. As such, data concatenation is only appropriate between conditions where the signal profile is consistent (e.g., same stimulation site on same day).

If it is expected that the artifacts differ across the compared datasets and concatenation is not possible, spatial filtering, such as SSP-SIR, allows the subsequent out-projection of both artifact subspaces. Although this approach might result in greater overall attenuation of the neuronal signals of interest, it ensures that the compared datasets suffer from the identical overcorrection.

### 5.6. Do not trust the default values

Another important consideration, particularly when using open-source toolboxes for performing analysis, is how to select the input parameters used by different cleaning methods. Often, the outcome of a given cleaning method will depend on the input parameters that control how the method performs (e.g., the lambda value for SOUND controls how much ‘noise’ is suppressed within the signal; the heuristic threshold levels in the TESA component selection function determine component classifications etc.). In most toolboxes, these parameters are assigned default values, often determined by how the cleaning method performed on a certain test data set. However, these default values are not always optimal for new data sets, which may have differing features (e.g., sampling rates, number of electrodes etc.) and artifact profiles (e.g., larger decay artifacts, TMS-evoked muscle activity etc.) to the original data set. When using default settings, it is important to carefully check the output of the cleaning method for unwanted or suboptimal behaviour (see recommendation 7 below). Using a range of values can also help to better understand the impact of a given input on cleaning performance.

On the other hand, the input settings should not be treated as free parameters that can be adjusted until the desired research outcome is reached. Although often challenging, one should always base the parameter settings on objective criteria directly related to the signal-processing step at hand where possible. The evaluation should also not be based on the same underlying assumptions as the adjusted cleaning method. For instance, if rejecting muscle artifacts with ICA that is based on finding maximally independent components, one could utilize the time-frequency criteria to find the number of independent components that need to be removed to sufficiently suppress the artifact (see section 7) (Mäki and Ilmoniemi, 2011; Mutanen et al., 2016). On the other hand, if one wishes to use the SOUND algorithm to clean noisy channels (Mutanen et al., 2018), one could look for the regularization parameter that is the best compromise between signal-to-noise ratio (SNR) and overcorrection. Because SOUND aims at maximizing SNR by cross validating EEG channels, the SNR estimate, used for determining the regularization parameter, could be determined, e.g., based on frequency analysis as in (Nikulin et al., 2011) and the overcorrection could be quantified, for example, using forward modelling as in (Salo et al., 2020).

### 5.7. Regularly visualise the data

A simple tip for helping design TMS-EEG analysis pipelines is to visualise the data after each cleaning step. Visualising the data serves two purposes. First, it allows for quick assessment of whether the artifacts targeted by the cleaning step have been adequately suppressed. Second, it facilitates detection of unwanted distortions that can be introduced to the data following cleaning. For example, the introduction of ringing or offsets before the stimulus pulse following filtering can indicate that there is a step in the data that is interacting with zero-phase filters. While this recommendation is subjective in nature and requires a level of familiarity with identifying artifacts and unwanted signal distortions, the importance of regularly visualising the data cannot be underestimated.

In addition to visualizing the data often during the preprocessing phase, it is recommended to visualize the data using various plot types. While the channel time courses might reveal, e.g. filtering artifacts like ringing, topographical plots are effective in highlighting contaminated channels. On the other hand, time-frequency transformation of the channel time courses can expose residual line noise, which introduces a sharp peak at 50 or 60 Hz, or TMS-evoked muscle artifacts, which results in a broad-band response immediately after the pulse.

## 6. The choice of cleaning steps matter: a practical example using the TESA toolbox

The above recommendations provide some broad guidance on designing TMS-EEG pipelines. However, there are still a number of choices that are required regarding which cleaning methods to use and in which specific order. An important question is whether these choices make any difference to the final TEP. If differing pipelines converge onto a unified TEP, then it does not matter which pipeline is used to clean TMS-EEG data. However, a growing number of studies have reported differences in TEP outcomes when using different cleaning methods to suppress TMS-evoked muscle and decay activity, including ICA, PCA, and detrending models (Casula et al., 2017; Hernandez-Pavon et al., 2012; Korhonen et al., 2011; Rogasch et al., 2014), suggesting the choice of cleaning method does matter. Furthermore, a recent study compared ‘default’ pipelines from different open-source TMS-EEG toolboxes (Bertazzoli et al., 2021). Three of these pipelines used similar cleaning methods (e.g., ICA) but in differing orders, and a fourth using SSP-SIR and SOUND. The study found that the final TEPs differed in nearly all pipeline comparisons, suggesting that the order in which cleaning steps are performed (and possibly the toolbox used) also matters, even when the same methods are applied.

Most of the above comparisons have used TMS-EEG data with complex artifact profiles, including TMS-evoked muscle activity, decay artifacts and, in some cases, time-locked blinks. It remains less clear whether different TMS-EEG cleaning pipelines also diverge on data with ‘simpler’ artifact profiles (e.g., decay artifacts with no TMS-evoked muscle activity). To test whether using different methods to suppress decay artifacts also results in differing TEPs, and to provide a practical example on how to use TESA to compare analysis pipelines, we compared TEPs following three different pipelines; one using FastICA, one using SOUND, and one using a detrending model. We kept the pipelines as consistent as possible except for the decay artifact cleaning step and used a data set with a well characterised artifact profile. We hypothesised that using different cleaning methods would result in different amplitudes and spatial distributions of the final TEP.

### 6.1 Methods

#### Data

The data used in this analysis were collected as part of a larger, unpublished study involving magnetic resonance imaging (MRI), resting-state EEG, task-based EEG, and TMS-EEG stimulating the left dorsolateral prefrontal cortex, superior frontal gyrus, posterior parietal cortex, and shoulder. All participants were healthy young individuals with no contraindications to MRI or TMS, and provided written informed consent prior to data collection. The protocol was approved by the Monash Human Research Ethics Committee (ID: 12723). For this analysis we used data from TMS-EEG when the superior frontal cortex was stimulated, and we selected individuals where transients from the TMS pulse had recovered within 10 ms, and who did not show TMS-evoked scalp muscle activity (n=14, 8 females, mean age = 33 years, range = 19-45 years). We deliberately avoided TMS-evoked muscle artifacts to ensure any TMS-evoked decay artifacts were not caused by the tail of muscle activity, or movement artifacts, thereby ensuring a well-characterised artifact profile within the data set.

Biphasic TMS pulses were delivered through a figure-of-eight coil connected to MagVenture X100 stimulator with MagOption (MagVenture). The coil position was determined using stereotaxic neuronavigation (BrainSight, Rogue Resolutions) targeting the left superior frontal gyrus (MNI coordinates: -23, 4, 63). The coil position and angle were adjusted so the centre of the coil was located over the nearest gyrus, and the coil handle ran perpendicular to the primary direction of the gyrus (i.e., to ensure maximal current flow in the gyrus). Stimulation intensity was determined based on the resting motor threshold from the left motor cortex adjusted for coil-to-cortex distance at the stimulation site using the formula described by Stokes and colleagues (Stokes et al., 2005). Stimulation was delivered at 120% of the adjusted threshold (mean = 77.2 ± 11% of maximum stimulator output) and 120 TMS pulses were delivered using an interstimulus interval jittered between 4-6 s. White noise was played through headphones throughout stimulation to help mask the TMS clicking noise, although most participants still perceived the click. EEG data were collected from 62 TMS-compatible sintered Ag-AgCl electrodes with a low profile, C-ring slit design (EASYCAP) arranged in standard 10-10 positions across the scalp. Data were online referenced to the FPZ electrode, with AFZ serving as the ground electrode. Electrode positions were digitised using the stereotaxic neuronavigation system. The EEG signals were amplified (1000x), filtered (DC-1,000 Hz), and recorded at a sampling rate of 10 kHz (Synamps RT amplifier, Compumedics). Electrode impedances were kept below 5 kΩ.

#### Cleaning pipelines

We compared TEPs following three different cleaning pipelines which were nearly identical except for the method used to suppress the TMS decay artifact. The pipelines were developed using EEGLAB (Delorme and Makeig, 2004) and TESA (Rogasch et al., 2017) on the Matlab platform. Topoplots were generated using FieldTrip (Oostenveld et al., 2011). Data were epoched around the TMS pulse (−1000 to 1000 ms), baseline corrected by subtracting the mean between -500 to -10 ms from all data points, and data around the TMS pulse was removed (−2 to 10 ms). Channels and trials showing excessive artifacts (e.g., persistent drifts, excessive muscle activity or line noise) were manually identified and removed. Removed channels were replaced using spherical interpolation, and the data were re-referenced to the average of all channels (note that missing channels were removed again prior to FastICA to ensure correct rank of the data).

At this point the common pipeline was separated into three parallel pipelines each using a different approach to suppress the TMS-evoked decay artifact. In pipeline 1, FastICA (Hyvarinen, 1999) was run on the data and components representing the decay artifact were automatically selected using heuristic rules in the *tesa_compselect.m* function (defined as components where the signal power in the data between 11-30 ms accounted for >8 times the signal power between 30-1000 ms). Components were manually checked and additional components removed if necessary (n=2). In pipeline 2, SOUND (Mutanen et al., 2020, 2018) was run on the data using individualised leadfield matrices for source estimation (generated using a 3-layer OpenMEEG boundary element model in the Brainstorm toolbox (Tadel et al., 2011)). In pipeline 3, the model proposed by Freche and colleagues (Freche et al., 2018) was fit to each trial and channel between 11-30 ms and then extrapolated to 11-500 ms using the same settings as the example in figure 12 of the original paper. The three electrodes with the most positive deflection and the three with the most negative deflections were used to define the model. The optimal model fits were then subtracted from the post-stimulation time window (11-500 ms) and the data were re-epoched to -500 to 500 ms. Following suppression of the decay artifacts, data from each pipeline were re-epoched to -500 to 500 ms to match the model pipeline. To assess the impact of each cleaning method on the data, TEPs from each pipeline were compared at this intermediate point in the cleaning pipelines.

Following the intermediate comparison, missing data from around the TMS pulse were interpolated using a cubic function (fit on 1 ms of data before and after the pulse), and the data were downsampled to 1000 Hz. The data were then band-pass (1-80 Hz) and band-stop (48-52 Hz) filtered using a zero-phase finite response filter (*pop_eegfiltnew.m*). FastICA was run on the data and components representing eye blinks, lateral eye movement, and persistent muscle activity were automatically selected and removed using heuristic rules in the *tesa_compselect.m* function (eye blinks: components where the weightings from FP1 and FP2 electrodes were >2.5 times other electrodes; lateral eye movement: where the weightings of F7 and F8 electrodes were +/- 2 times other electrodes respectively; muscle: components where the slope of the background power spectra in log-log scaling was > -0.31 [fit between 7-70 Hz excluding 48-52 Hz] (Fitzgibbon et al., 2016)). To help standardise comparisons between pipelines, the component selections were not manually adjusted following this step. Missing channels were interpolated again using spherical interpolation and the final TEPs were compared between the pipelines.

#### Analysis and statistics

For comparisons at the intermediate and end timepoints, we first averaged the data across trials to calculate TEPs. To assess differences in amplitude between pipeline outcomes, we calculated the global mean field amplitude (GMFA) by taking the standard deviation across all channels for each timepoint. For statistical comparisons, we extracted data following the TMS pulse (11-250 ms), and downsampled the data to 250 Hz resulting in 60 timepoints. Paired-sample t-tests were used to compare GMFA values between each combination of pipelines for each timepoint. To assess topographical similarities between pipeline outcomes, we first used Pearson’s correlations to compare TEP topographies between each pipeline combination for every timepoint and individual. Correlation coefficients were then converted to z scores using Fisher’s r to z conversion and one-sample t-tests were used to assess whether z scores differed from zero for each timepoint. For each set of comparisons, p<0.05 was considered significant and we used the Benjamini-Hochberg method to correct the false discovery rate across all 60 timepoints.

#### Data and code availability

Data used in these analyses are available from: https://doi.org/10.26180/18805994.v4

Code is available from: https://github.com/nigelrogasch/TMS-EEG-cleaning-pipelines

### 6.2 Results and discussion

Following the initial cleaning steps, on average 15.6 ± 9 trials (range = 2-31) and 2.6 ± 2 electrodes (range = 1-6) were removed from the data. Figure 2 shows group mean TEP butterfly plots for the raw data (following epoching, baseline correction, removal of trials/electrodes, and average re-referencing), the data following suppression of the decay artifact using each different processing method (intermediate comparison point), and the final TEPs at the end of each pipeline. The electrode closest to the coil (FC1) is highlighted in each plot and shows a considerable decay artifact prior to processing.

**Figure 2:**
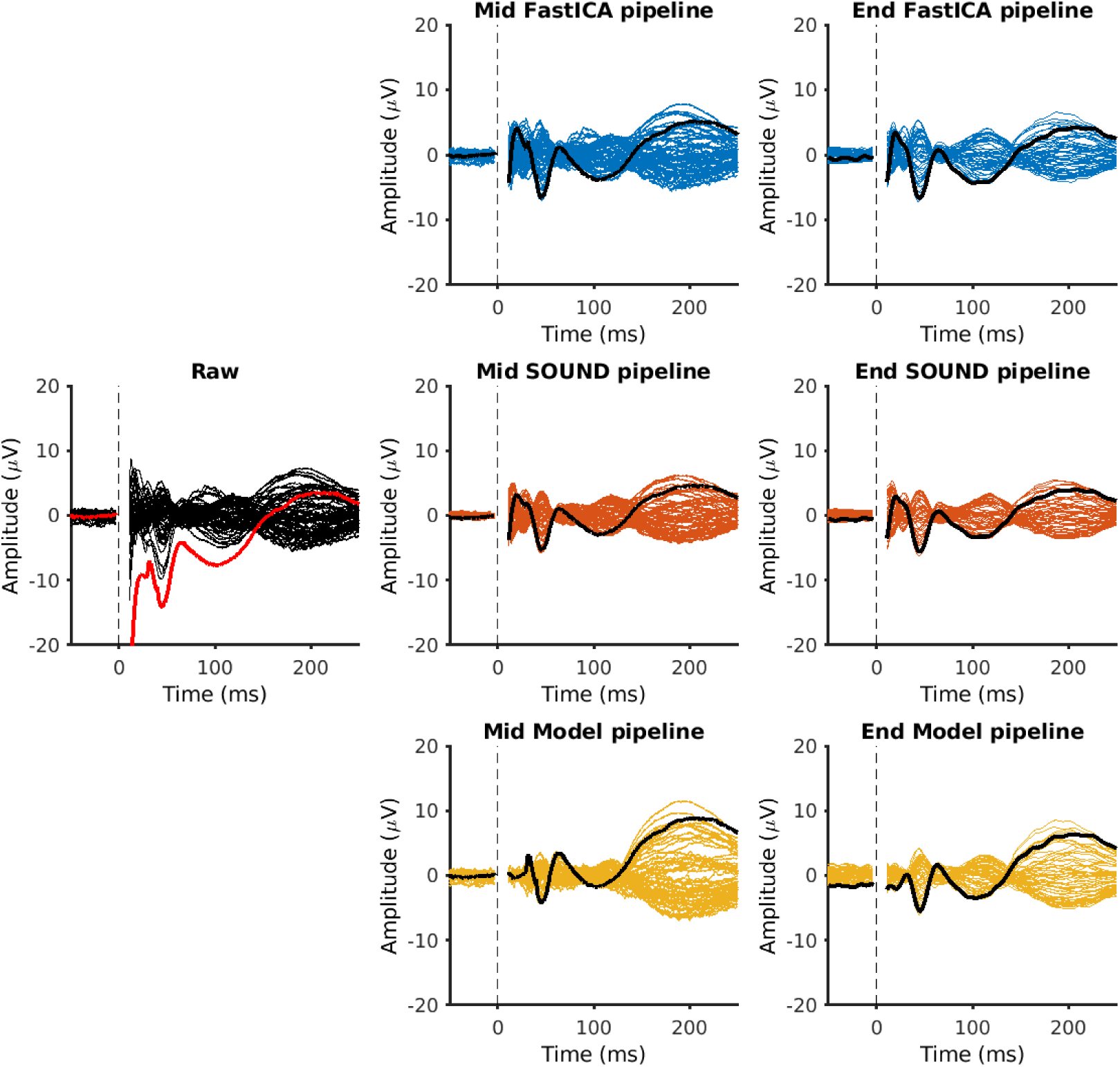
butterfly plots of group average TEPs from all electrodes at different stages of cleaning. TEPs in the left column are following the initial cleaning steps (epoching, baseline correction, trial/electrode removal, average re-reference), TEPs in the middle column are immediately following the three different decay suppression methods (FastICA in the top row, SOUND in the middle row, and the model in the bottom row), and TEPs in the right column are at the end of the three different pipelines. Data from the electrode closest to the coil (FC1) are highlighted in each plot (red in left column, black in middle and right columns).

At the intermediate comparison point (after removing the decay artifact), all three pipelines showed differences in GMFA and topographies (fig. 3). FastICA and SOUND pipelines resulted in similar GMFA patterns, however the amplitude was higher across nearly all timepoints following FastICA. The topographies were similar following FastICA and SOUND pipelines, with correlations >0.6 from 13 ms and >0.8 from 38 ms. In contrast, the model pipeline resulted in lower amplitudes within the first 120 ms and higher amplitudes after 130 ms compared to both FastICA and SOUND. Furthermore, the topography correlations between the model pipeline and both FastICA and SOUND were consistently lower than comparisons between the other two pipelines, especially within the first 45 ms following the pulse. Together, these findings highlight the differences in approach for suppressing the TMS-evoked decay artifact between methods.

**Figure 3:**
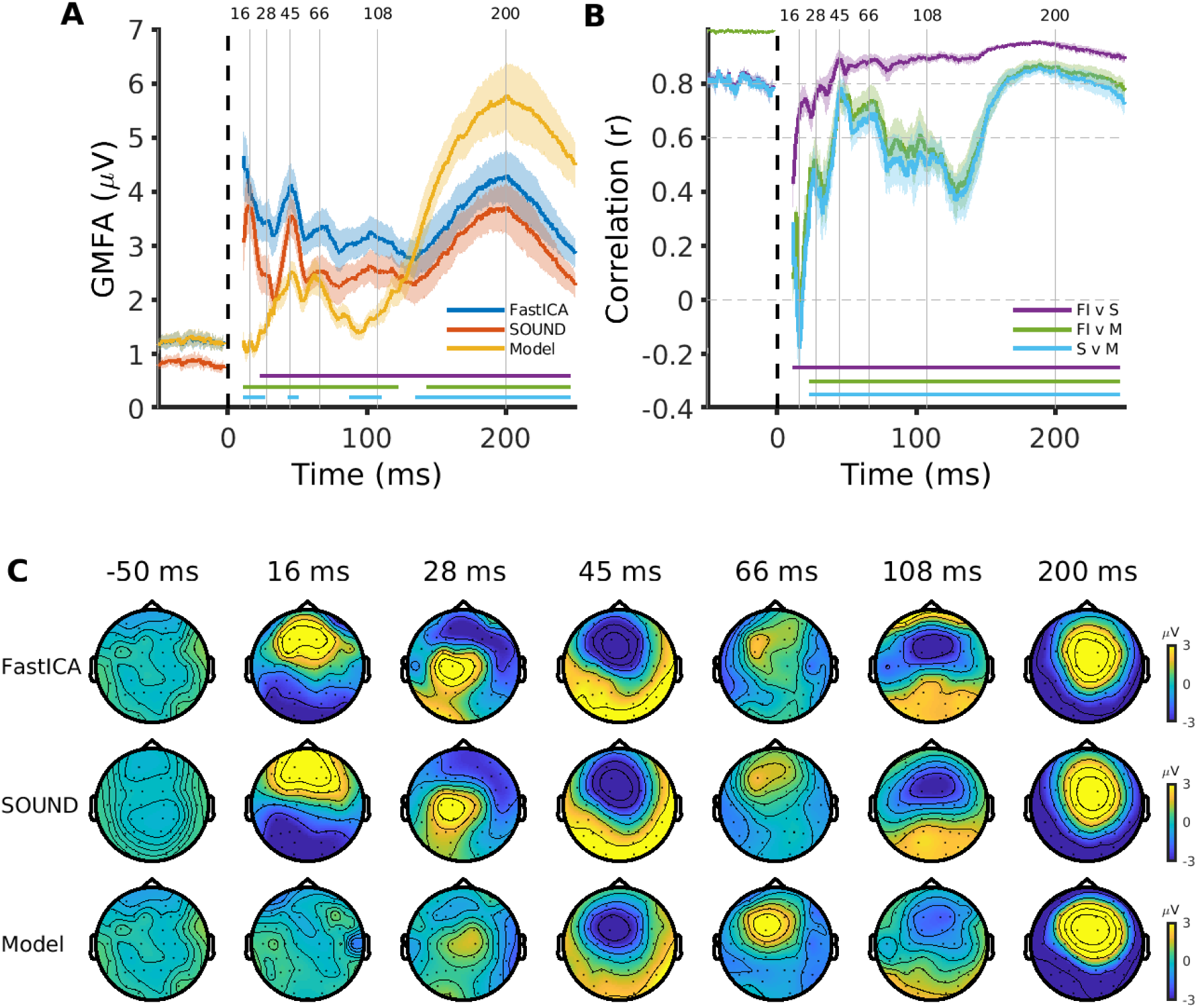
TEP comparisons between pipelines following the decay artifact suppression step. A) Global mean field amplitude (GMFA) of TEPs following each cleaning step. Solid lines (blue, orange, yellow) represent the group mean and shaded lines the standard error. Horizontal lines represent timepoints showing statistically significant differences between pipelines (p<0.05, FDR corrected; purple = FastICA vs SOUND [FI v S]; green = FastICA vs model [FI v M]; and light blue = SOUND vs model [S v M]). Dotted vertical line indicates the timing of the TMS pulse and grey vertical lines indicate the timing of peaks in the data. B) Pearson’s correlations between scalp maps from different pipelines across timepoints. Horizontal lines indicate correlations greater than 0 (p<0.05, FDR corrected). C) Topoplots showing the spatial distribution of EEG amplitudes across the scalp at peaks indicated in A and B following each pipeline.

At the end of the pipelines, most of the differences between FastICA and SOUND pipelines were removed, with both pipelines showing similar GMFA and only small differences in amplitude between 27-39 ms and 119-123 ms (fig. 4). Furthermore, both pipelines resulted in highly similar topographies across nearly all timepoints (correlations >0.6 from 12 ms; >0.8 from 34 ms). The resolution between the pipelines likely reflects the differences in approaches between the SOUND and FastICA pipelines. SOUND is designed to suppress biophysically implausible noise from the signal, which includes decay artifacts but also muscle artifacts and other noise sources, leaving fewer artifacts to suppress with the later FastICA step. In the FastICA pipeline, only components representing the decay artifact are removed in the first FastICA step, with muscle and other noise suppressed in the later FastICA step. In support of this interpretation, less variance was removed from the later FastICA step in the SOUND pipeline (mean = 9.1 ± 7%) compared to the FastICA pipeline (mean = 28.1 ± 16%; p = 8.1×10^-4^, paired-sample t-test). The model pipeline showed lower amplitudes at early timepoints (∼11-30 ms), higher amplitudes at later timepoints (∼140-190 ms), and lower topographical correlations across all timepoints (<0.6 until 40 ms, <0.8 until 160 ms) compared with the FastICA and SOUND pipelines. Of note, the model pipeline also showed higher amplitudes during the baseline period, possibly reflecting additional artifacts caused by band-pass filtering.

**Figure 4:**
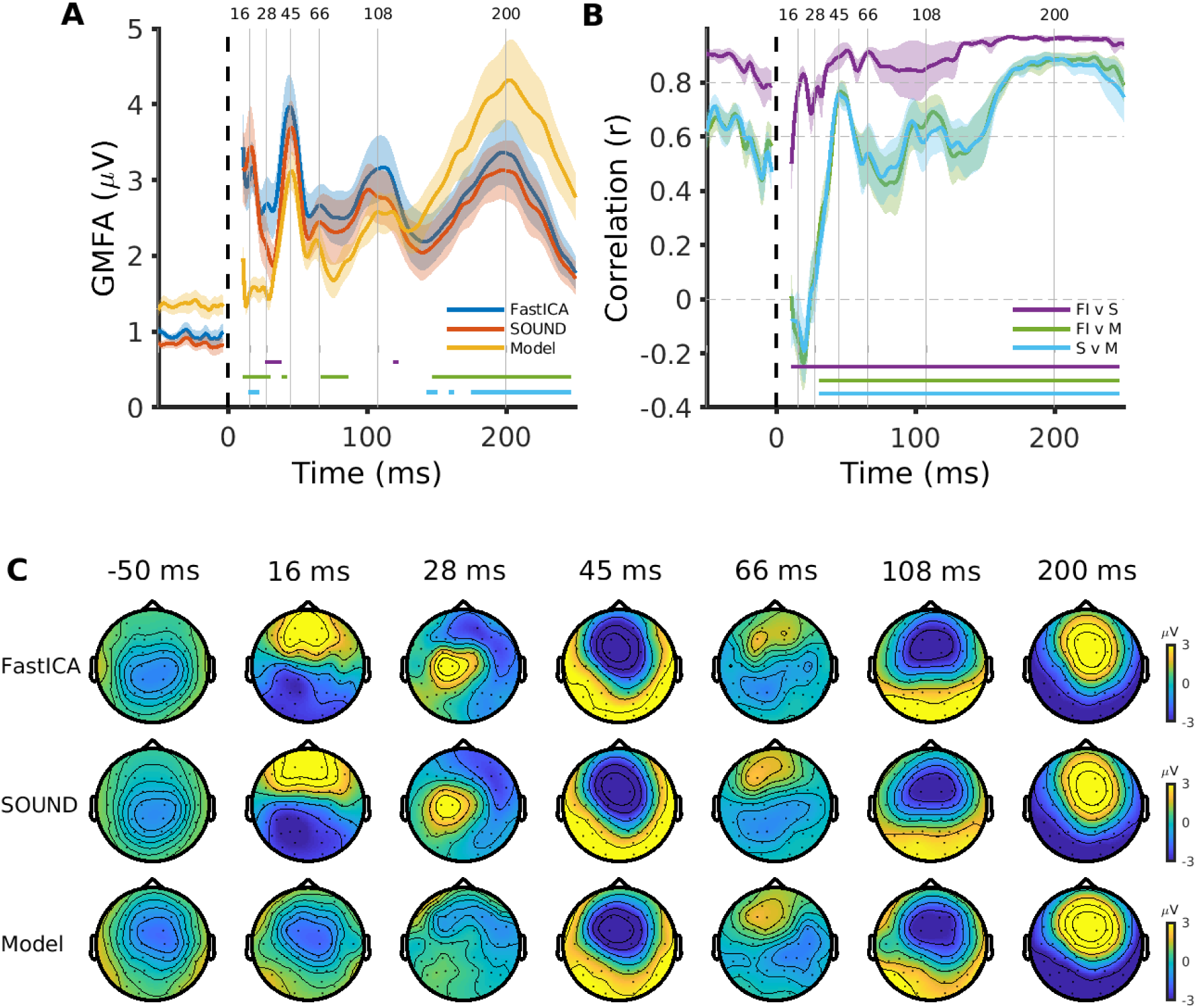
TEP comparisons between pipelines at the end of cleaning. A) Global mean field amplitude (GMFA) of TEPs following each pipeline. Solid lines (blue, orange, yellow) represent the group mean and shaded lines the standard error. Horizontal lines represent timepoints showing statistically significant differences between pipelines (p<0.05, FDR corrected; purple = FastICA vs SOUND [FI v S]; green = FastICA vs model [FI v M]; and light blue = SOUND vs model [S v M]). Dotted vertical line indicates the timing of the TMS pulse and grey vertical lines indicate the timing of peaks in the data. B) Pearson’s correlations between scalp maps from different pipelines across timepoints. Horizontal lines indicate correlations greater than 0 (p<0.05, FDR corrected). C) Topoplots showing the spatial distribution of EEG amplitudes across the scalp at peaks indicated in A and B following each pipeline.

To further investigate the discrepancies between the model pipeline and other pipelines, we plotted the model fits compared to the raw data for one participant (fig. 5A). These plots suggest the model was overfitting the shape of the early decay activity, and instead fitting the shape of the early TEP components. Furthermore, the extrapolated fit showed a poor match with later timepoints, explaining the larger amplitudes observed after 140 ms in fig. 4. To overcome these issues, we reran the model pipeline using a simplified model (from figure 10 in the original paper; model defined on one electrode with most positive deflection and one with most negative deflection) fit over a longer period (11-70 ms), which resulted in a model better capturing the decay artifact without fitting the early TEP peaks (fig. 5B). When comparing outcomes using this approach, the model pipeline TEPs looked more similar with the FastICA and SOUND pipelines (fig.6). There were small differences in the amplitude of the very early timepoints (11-16 ms) between the other pipelines which did not reach statistical significance. Furthermore, the topographical correlations were slightly lower for the first 16 ms and for the TEP peaks at 28 and 66 ms, however the correlations with other pipelines were generally high across time (>0.6 ms from 14 ms with both pipelines). These findings further highlight the importance of selecting appropriate parameters for a given data set when designing pipelines and visualising the outcomes to ensure cleaning steps are not introducing spurious signals to the data.

**Figure 5:**
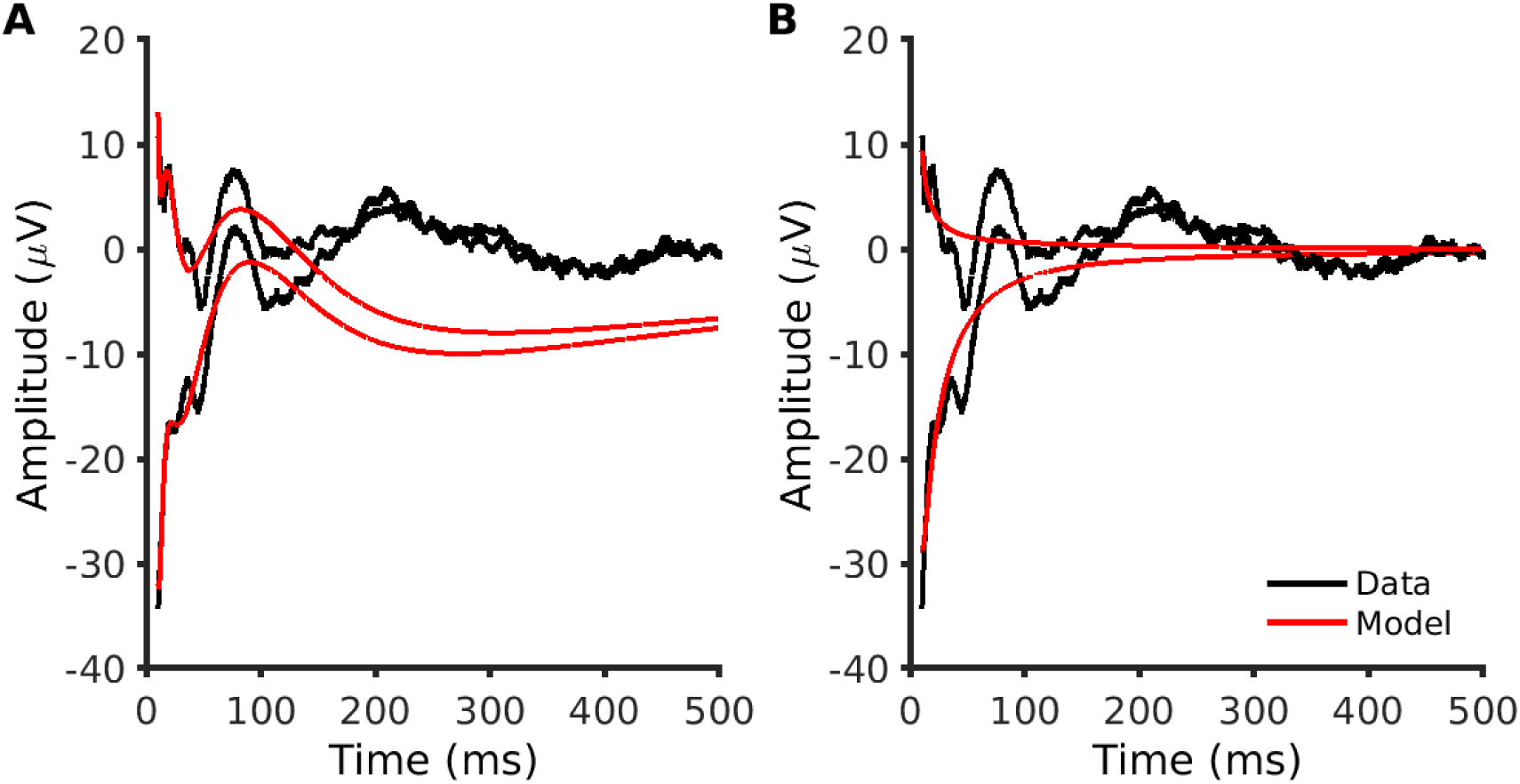
comparisons of TEPs and model fits from a representative individual. A) TEPs from two electrodes (FC1 and F3; black lines) and mean model fits (red lines) using the initial model. B) TEPs from the same two electrodes in A and mean model fits using an alternative model. The alternative model better captures the decay artifact without overfitting early TEP peaks.

**Figure 6:**
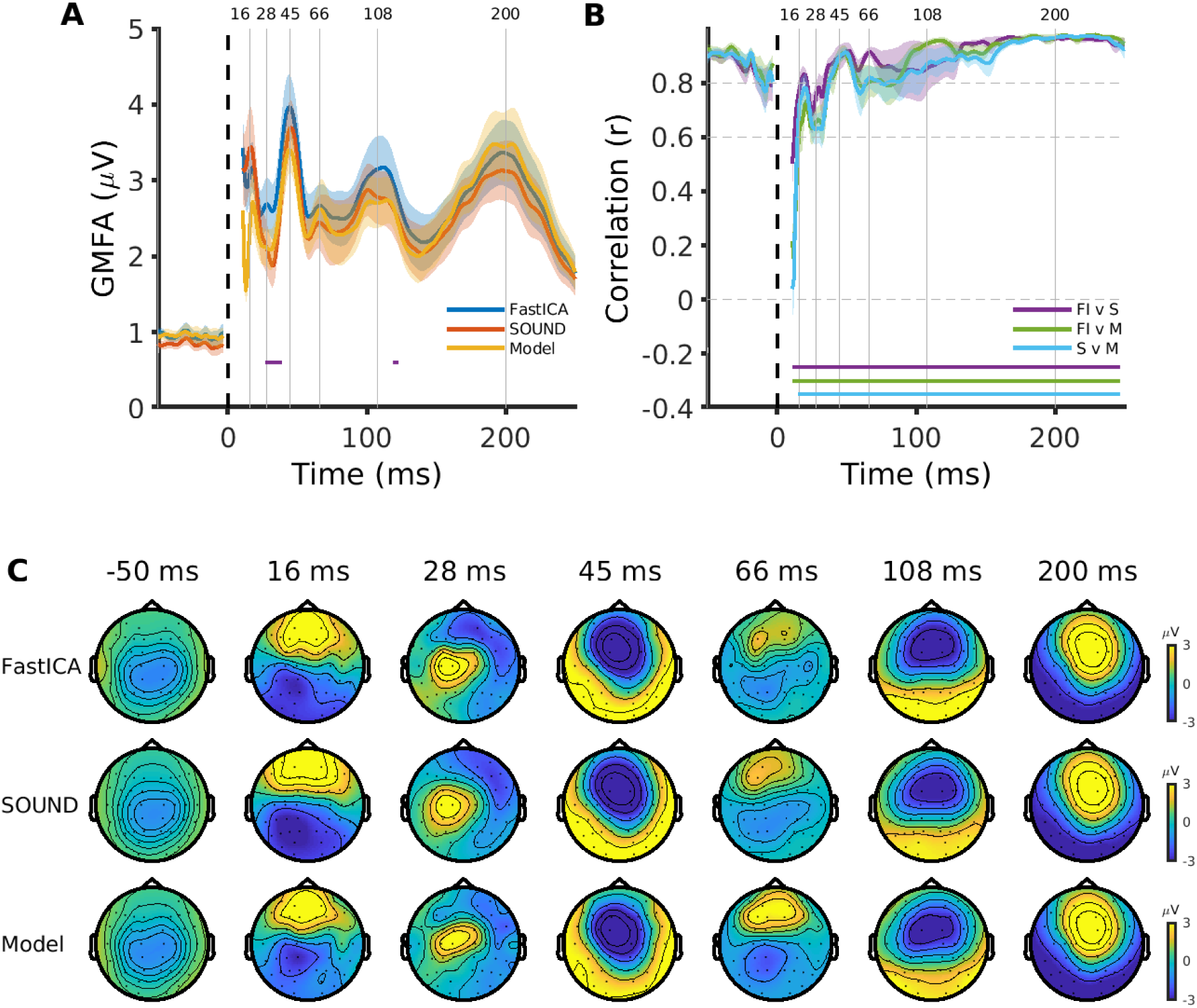
TEP comparisons between pipelines at the end of cleaning using the alternative model pipeline. A) Global mean field amplitude (GMFA) of TEPs following each pipeline. Solid lines (blue, orange, yellow) represent the group mean and shaded lines the standard error. Horizontal lines represent timepoints showing statistically significant differences between pipelines (p<0.05, FDR corrected; purple = FastICA vs SOUND [FI v S]; green = FastICA vs model [FI v M]; and light blue = SOUND vs model [S v M]). Dotted vertical line indicates the timing of the TMS pulse and grey vertical lines indicate the timing of peaks in the data. B) Pearson’s correlations between scalp maps from different pipelines across timepoints. Horizontal lines indicate correlations greater than 0 (p<0.05, FDR corrected). C) Topoplots showing the spatial distribution of EEG amplitudes across the scalp at peaks indicated in A and B following each pipeline.

Another possible source of variation between pipelines is FastICA. As the starting parameters of FastICA are seeded randomly, repeated runs can lead to small variations in the resulting components. Furthermore, additional variability can be introduced by inconsistent component removal between runs. To assess how consistent outcomes were within pipelines, we repeated the FastICA steps for each pipeline an additional two times. Of note, we used heuristic rules to help standardise component selection between pipelines and minimise variability. TEPs from the same pipeline were highly similar for all three pipelines evaluated (fig. 7), suggesting that the small differences between pipelines observed in figure 6 were likely the result of the method used to remove the decay artifact as opposed to variability resulting from FastICA. These findings also demonstrate the high reproducibility of TEPs within a given pipeline (i.e., TEP outcomes are very consistent across repeated iterations of the same pipeline despite potential variability in FastICA decomposition and selection).

**Figure 7:**
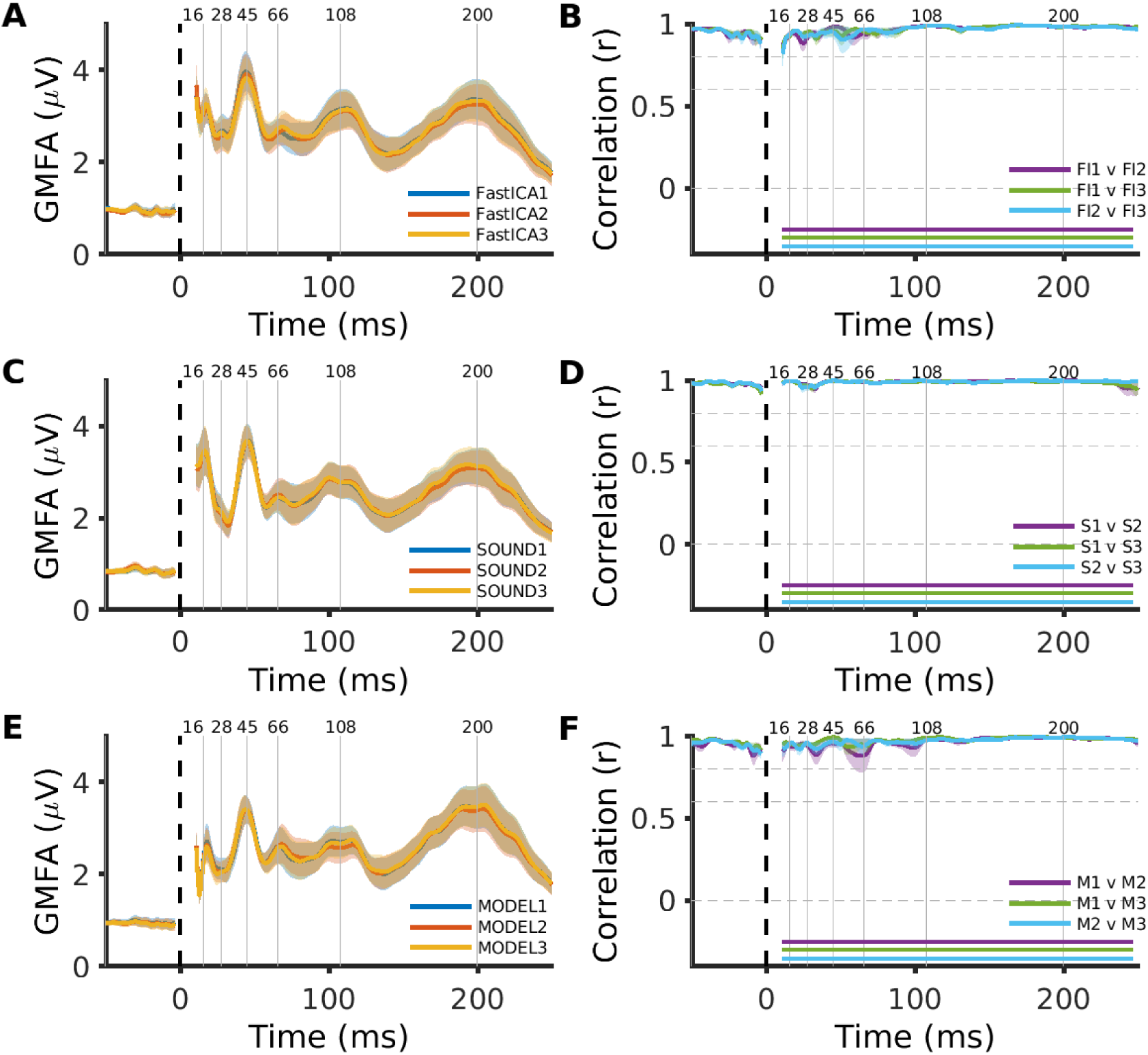
TEP comparisons following repeated analysis using the same pipeline. Left column: global mean field amplitude (GMFA) of TEPs following three repeats of each pipeline. Solid lines (blue, orange, yellow) represent the group mean and shaded lines the standard error. Right column: Pearson’s correlations between scalp maps from repeats within pipelines across timepoints.

Finally, we assessed whether changing the method used for correcting noisy channels altered TEP outcomes. The SOUND algorithm was designed to correct noisy channels automatically, and therefore does not require manual removal and interpolation of channels, as was performed in the above pipelines. Therefore, we compared TEPs following the SOUND pipeline used in the above analyses to TEPs following a pipeline where noisy channels were not removed prior to SOUND. TEPs were very similar regardless of whether noisy channels were removed manually or corrected using the SOUND algorithm (fig. 8), suggesting the choice of channel correction methods does not have a large impact on TEP outcomes for these pipelines.

**Figure 8:**
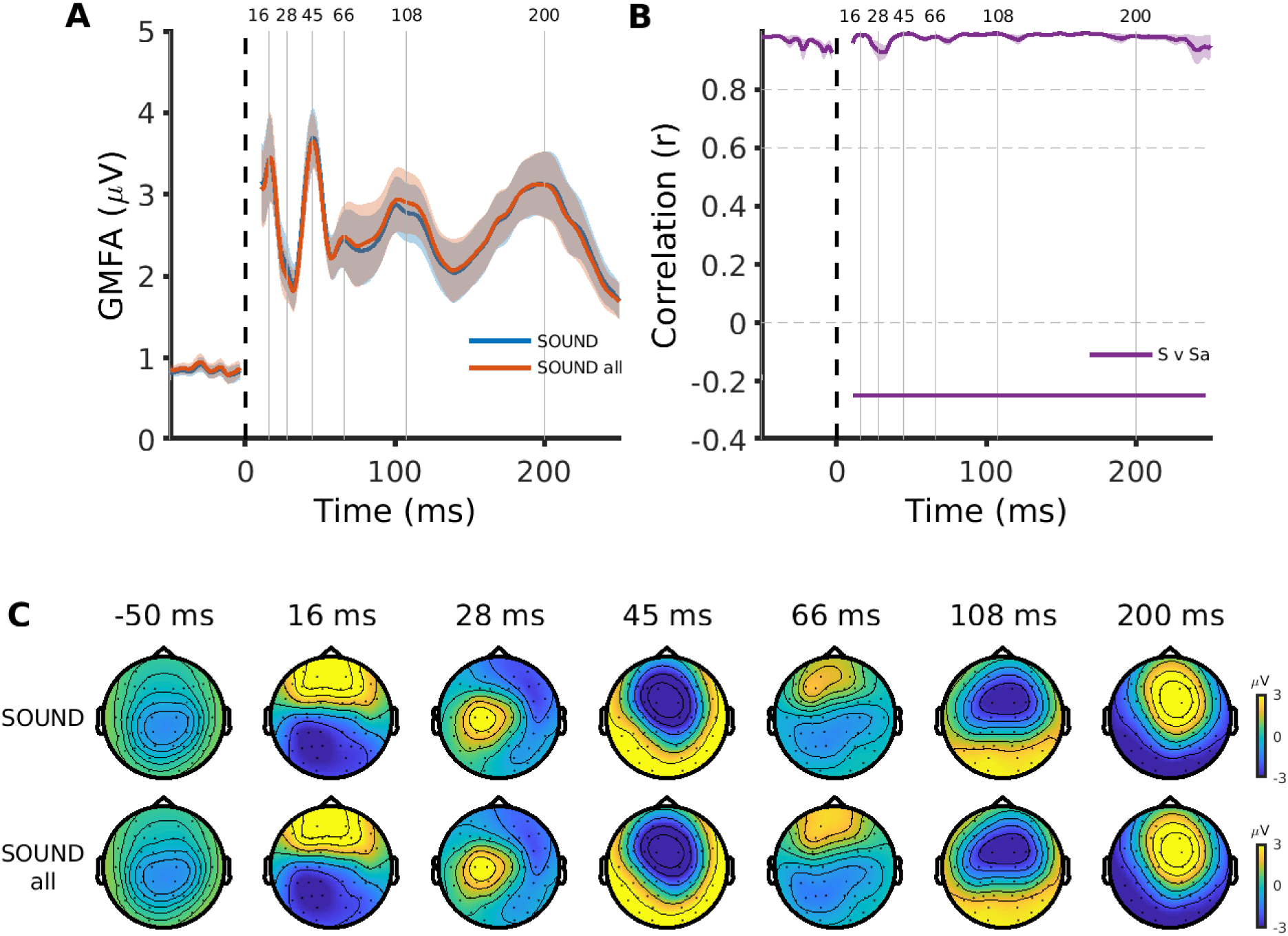
TEP comparisons between pipelines following different methods for correcting noisy channels. A) Global mean field amplitude (GMFA) of TEPs following each cleaning pipeline. Solid lines (blue, orange) represent the group mean and shaded lines the standard error. Dotted vertical line indicates the timing of the TMS pulse and grey vertical lines indicate the timing of peaks in the data. B) Pearson’s correlations between scalp maps from different pipelines across timepoints. Horizontal lines indicate correlations greater than 0 (p<0.05, FDR corrected). C) Topoplots showing the spatial distribution of EEG amplitudes across the scalp at peaks indicated in A and B following each pipeline.

To summarise, we found that changing a single step in the TMS-EEG cleaning pipeline used to suppress decay artifacts resulted in TEPs that were broadly similar, but showed small differences in amplitude and topography, particularly for early latency timepoints. In contrast, TEPs were highly consistent following repeated runs of the same pipeline, and when using different methods for correcting noisy channels. Our findings are consistent with other recent reports showing differences in TEPs between pipelines from different TMS-EEG analysis toolboxes (Bertazzoli et al., 2021), and different approaches for suppressing decay artifacts (Casula et al., 2017). In contrast to previous studies, we carefully selected data with similar artifact profiles (decay, but not TMS-evoked muscle artifacts), only varied one step in the cleaning pipeline, and compared within-pipeline consistency to better isolate the cause of variability in pipeline outcomes. Our findings also reinforce the advice from the previous section - the settings used to generate the model for fitting the decay artifact had a large impact on the final result, and this discrepancy was detected by visualising the data. Together, these findings highlight the challenges inherent in cleaning TMS-EEG data - the choice of cleaning steps matter, even for data with minimal artifact profiles.

## 7. Which analysis pipelines should I use? The challenges of validation

In the above analysis, we showed that changing a single step in an analysis pipeline can result in different TEP outcomes, particularly for early TEPs. Furthermore, others have shown similar results when comparing similar pipelines from different toolboxes (Bertazzoli et al., 2021). So which analysis pipeline is the ‘best’ one to use? We often see papers using a certain pipeline or method and citing the pipeline is ‘validated’, thereby implying that the resulting ‘cleaned’ TEPs are an accurate representation of the EEG activity evoked by TMS. For example, we have seen this often for the two-step ICA approach introduced by our team (Rogasch et al., 2014). However, truly validating a cleaning pipeline is extremely challenging, mainly because the ‘ground truth’ (i.e., the signal we are trying to uncover) is usually unknown. To our knowledge there are no TMS-EEG pipelines that we would consider adequately validated against a known ground truth to enable confident recommendation over other pipelines. Indeed, generating a ground truth for benchmarking against is a considerable challenge. This problem is not unique to TMS-EEG, and is shared by many other neuroimaging methods including EEG, magnetoencephalography, and functional MRI, which can all suffer from low signal-to-noise ratios. There are several approaches which could be used to generate benchmarking data sets for pipeline validation. One approach is to optimise online methods and experimental designs to minimise artifacts during data collection as much as possible. This approach maximises signal-to-noise ratios and facilitates the use of minimal cleaning pipelines. It may be possible to then introduce known artifacts and test the ability of different pipelines for reproducing the ‘clean’ data set. For example, one could remove the noise masking, thereby reintroducing auditory-evoked potentials (Rocchi et al., 2021). There is also anecdotal evidence that some amplifiers show larger decay artifacts than others – collecting data on both systems with the same individuals using the same EEG caps could provide a data set for validating decay artifact removal. Another approach is to simulate known biologically-plausible signals using biophysical models (Breakspear, 2017). There are several models capable of capturing key features of EEG signals (Cona et al., 2011; Sanz-Leon et al., 2018), however additional work on modelling the various artifact sources such as TMS-evoked muscle and decay artifacts is required. Until benchmarking data sets are developed, and pipelines validated, it is difficult to recommend one specific TMS-EEG cleaning pipeline.

## 8. Recommendations for cleaning TMS-EEG data

In the previous sections, we have shown that there is an enormous combination of different possible cleaning pipelines for TMS-EEG data and changing even a single step within an analysis pipeline can alter the resulting TEPs. Furthermore, we have argued that there is currently no validated ‘best’ pipeline for analysing TMS-EEG data and that we may even need different analysis pipelines for different TMS-EEG data sets. Given these challenges, what can we do to minimise variability in the outcome from TMS-EEG cleaning pipelines, both within and between studies?

### 8.1. Analyse data using multiple pipelines

The main strategy we propose to minimise analysis variability within studies is to analyse TMS-EEG data using multiple pipelines. The so-called ‘multiverse analysis’ approach has been suggested for other methods and fields facing analytical variability including functional MRI (Botvinik-Nezer et al., 2020) and psychology (Silberzahn et al., 2018). The main goal of this approach is to confirm an effect or lack of an effect (e.g., a difference in TEP characteristics between conditions, or groups) is not due to variability caused by a specific analysis pipeline. While a true multiverse analysis requires numerous different analysis approaches and a meta-analysis of the results, a practical alternative is to compare results from 2-3 analysis pipelines. This approach has been adopted by several recent TMS-EEG studies (e.g., Biabani et al., 2019; Rogasch et al., 2020) and was also advocated by a recent study comparing TMS-EEG analysis toolboxes (Bertazzoli et al., 2021). Importantly, a multiple pipeline approach allows assessment of how robust a given finding is to changes in the preprocessing methods. While the specificity of a result to a given pipeline does not invalidate the finding per se, it does provide important insight into potential factors underlying differences in TEPs.

TESA is specifically designed to allow the combination and comparison of different TMS-EEG cleaning pipelines. To facilitate the comparison of multiple analysis pipelines, we have made all data and code available from the analysis used in the current study to provide examples on how this can be achieved using TESA. We have also created a pipeline repository within TESA which provides example scripts for different published pipelines which can be fully implemented within the TESA and EEGLAB framework. In the first iteration, we provide three pipelines using contrasting approaches: a minimal cleaning pipeline appropriate for data where TMS-evoked muscle and decay artifacts have been avoided online, and two pipelines designed for data containing TMS-evoked muscle/decay artifacts; one using the two-step ICA approach and one using SOUND and SSP-SIR. We expect this list of pipelines will grow as more approaches become available. Users can adapt these template pipelines to suit their own data. Given the open-source availability of different TMS-EEG analysis pipelines, we recommend the multiple pipeline approach becomes standard practice within the field until pipeline validation is improved.

### 8.2. Share data and code

Another practice we advocate is sharing the data and analysis code used within TMS-EEG publications (where ethically possible). Sharing data and code has become common in a number of neuroimaging fields, in particular MRI (Poldrack and Gorgolewski, 2014), and has numerous advantages. First, sharing data and code enhances reproducibility and transparency by enabling other researchers to directly replicate the results from published studies and makes it easier for these analysis approaches and cleaning pipelines to be adopted in new studies. Second, the availability of open TMS-EEG data creates a test bed for new analysis methods and pipelines. As new cleaning and analysis methods are developed, it will become increasingly important to test how these methods generalise across different data sets collected by different laboratories. Finally, access to TMS-EEG data will also allow more direct comparisons between TEPs collected by different groups. Such comparisons are important, especially given the current debates regarding the reproducibility and standardisation of TMS-EEG research (Belardinelli et al., 2019; Siebner et al., 2019), and will greatly assist in identifying the salient features driving variability in TEPs between studies.

While ideologically appealing, there are several challenges which accompany data sharing. The most pressing is ethical concerns regarding the potential for identifying individuals from their data. While this issue is more salient for methods like MRI where an individual’s facial features can be reconstructed from anatomical scans if not ‘defaced’ (Schwarz et al., 2019), there is evidence that TMS-evoked potentials are also highly specific to individuals using methods known as ‘fingerprinting’ (Ozdemir et al., 2021). Although extremely unlikely, this does raise the possibility that someone could be identified from an open-access data set based on their TEPs. Another important ethical consideration is the ability of participants to consent to how and by whom their data is used. Data sharing and ownership laws differ between jurisdictions and can create barriers (both real and perceived) for making data publicly available (White et al., 2020). However, many of these barriers can be overcome by adopting appropriate consenting procedures which clearly outline the proposed data sharing practices. Furthermore, there are a variety of ways in which data can be shared, ranging from publicly accessible data repositories to more restrictive approaches which require data sharing arrangements between the data host and applicant, thereby providing increased protection and control over who has access to data. Finally, a practical consideration is that the high sampling rates used to collect TMS-EEG data result in large data files (>1 GB per file) which can make data storage and sharing challenging. There are now numerous platforms for sharing scientific data (e.g., open science framework), some specifically designed for neuroimaging data sets which can accommodate large data files (e.g., OpenNeuro) (Markiewicz et al., 2021). Furthermore, platforms like OpenNeuro require standardised data formats such as Brain Imaging Data Structure (BIDS) (Gorgolewski et al., 2016), increasing data quality and transparency. Given the potential benefits for improving reproducibility and method development, we recommend data and code sharing become standard practice within the TMS-EEG field.

### 8.3. Choose the right tool for the job

Different preprocessing steps are designed for specific purposes, and thus, are based on different theoretical premises. Therefore, it is fundamental to understand at least the basic underlying assumptions of the applied methods to critically evaluate whether the applied signal processing tool is suitable for the data and artifact at hand. Failing to do so might lead to misinterpretations of the result. An excellent practical example was presented in this paper; when following immediately after the decay-artifact removal, SOUND seemingly attenuated the EEG signals the most. The reason for this, however, was that SOUND was never tailored for removing only decay artifacts. Instead, SOUND probably removed other signal components that were unlikely to originate from intracranial dipolar sources. On the contrary, we customized the FastICA and the model-fitting approaches for specifically removing the decay artifact. Accordingly, when ICA was finally used to clean the remaining artifacts, other than the decay, both the SOUND and ICA pipelines produced much more similar results.

A researcher should also always consider their research question and in its context when choosing the steps to include in a preprocessing pipeline. When TMS-evoked scalp muscle activity is present in the data, complicated preprocessing pipelines are required for quantifying the early responses to TMS. However, every added level of complexity in the data analysis increases the risk of introducing filtering artifacts that can be difficult to notice. Hence, even in the presence of prominent artifacts, it might be more beneficial to leave that artifact untouched if it does not overlap in time or frequency with the signals of interest. For instance, even large-amplitude muscle artifacts might not hinder the analysis of the N100 component because muscle responses are characteristically short-lived. For such a study, a simple temporal rejection and interpolation might be a much safer choice than ICA, which assumes that the data consists of statistically independent components. Ultimately, we recommend: 1) choosing analysis approaches that require the fewest assumptions wherever possible; and 2) appropriately acknowledging the limitations inherent in using complicated pipelines to clean highly artifactual data.

## 9. Conclusions

Retrieving neural activity from EEG data following a TMS pulse remains a challenge due to the unique artifacts resulting from the combination of techniques. While minimising these artifacts online by appropriate experimental design and careful preparation during data collection is essential, artifacts are difficult to avoid in all TMS-EEG experiments. Suppressing artifacts using offline cleaning methods is therefore an important component of TMS-EEG experimentation. The availability of methods for designing and implementing TMS-EEG cleaning pipelines is improving thanks to open science practices, like sharing data and code, and the development of specialised open-source toolboxes such as TESA. However, the choice of methods and the order in which they are implemented can alter the final TEP, even for data with a minimal artifact profile, as demonstrated by our current findings. The variability in TEP outcome following cleaning has important implications for comparing TMS-EEG results between different research groups, especially given the wide variety of online and offline artifact suppression approaches currently in use. The next challenge for offline approaches is designing robust experimental and simulation methods, which will allow validation of TMS-EEG cleaning pipelines against known ground-truths. Until then, it is recommended that researchers compare results from TMS-EEG studies using multiple cleaning pipelines to ensure findings are not related to the choice of cleaning methods. The continuing improvement and validation of offline cleaning methods will play an essential role in ensuring TMS-EEG reaches its full potential as a method for exploring the function of neural circuits in both basic and clinical research.

## Abbreviations

EEG: electroencephalography
TMS: transcranial magnetic stimulation
TEP: TMS-evoked potential
MEP: motor-evoked potential
ICA: independent component analysis
PCA: principal component analysis
SOUND: source-estimate-utilising noise-discarding
SSP-SIR: signal-space projection source-informed reconstruction
GUI: graphical user interface
ERP: event-related potentials
SNR: signal-to-noise ratio
MRI: magnetic resonance imaging
GMFA: global mean field amplitude
BIDS: Brain Imaging Data Structure

## References

Atluri, S., Frehlich, M., Mei, Y., Garcia Dominguez, L., Rogasch, N.C., Wong, W., Daskalakis, Z.J., Farzan, F., 2016. TMSEEG: A MATLAB-Based Graphical User Interface for Processing Electrophysiological Signals during Transcranial Magnetic Stimulation. Front. Neural Circuits 10, 78. https://doi.org/10.3389/fncir.2016.00078

Barker, A.T., Jalinous, R., Freeston, I.L., 1985. Non-invasive magnetic stimulation of human motor cortex. Lancet Lond. Engl. 1, 1106–1107. https://doi.org/10.1016/s0140-6736(85)92413-4

Belardinelli, P., Biabani, M., Blumberger, D.M., Bortoletto, M., Casarotto, S., David, O., Desideri, D., Etkin, A., Ferrarelli, F., Fitzgerald, P.B., Fornito, A., Gordon, P.C., Gosseries, O., Harquel, S., Julkunen, P., Keller, C.J., Kimiskidis, V.K., Lioumis, P., Miniussi, C., Rosanova, M., Rossi, S., Sarasso, S., Wu, W., Zrenner, C., Daskalakis, Z.J., Rogasch, N.C., Massimini, M., Ziemann, U., Ilmoniemi, R.J., 2019. Reproducibility in TMS-EEG studies: A call for data sharing, standard procedures and effective experimental control. Brain Stimulat. 12, 787–790. https://doi.org/10.1016/j.brs.2019.01.010

Bender, S., Basseler, K., Sebastian, I., Resch, F., Kammer, T., Oelkers-Ax, R., Weisbrod, M., 2005. Electroencephalographic response to transcranial magnetic stimulation in children: Evidence for giant inhibitory potentials. Ann. Neurol. 58, 58–67. https://doi.org/10.1002/ana.20521

Bertazzoli, G., Esposito, R., Mutanen, T.P., Ferrari, C., Ilmoniemi, R.J., Miniussi, C., Bortoletto, M., 2021. The impact of artifact removal approaches on TMS-EEG signal. NeuroImage 239, 118272. https://doi.org/10.1016/j.neuroimage.2021.118272

Biabani, M., Fornito, A., Coxon, J.P., Fulcher, B.D., Rogasch, N.C., 2021. The correspondence between EMG and EEG measures of changes in cortical excitability following transcranial magnetic stimulation. J. Physiol. https://doi.org/10.1113/JP280966

Biabani, M., Fornito, A., Mutanen, T.P., Morrow, J., Rogasch, N.C., 2019. Characterizing and minimizing the contribution of sensory inputs to TMS-evoked potentials. Brain Stimul. Basic Transl. Clin. Res. Neuromodulation 12, 1537–1552. https://doi.org/10.1016/j.brs.2019.07.009

Bonato, C., Miniussi, C., Rossini, P.M., 2006. Transcranial magnetic stimulation and cortical evoked potentials: a TMS/EEG co-registration study. Clin. Neurophysiol. Off. J. Int. Fed. Clin. Neurophysiol. 117, 1699–1707. https://doi.org/10.1016/j.clinph.2006.05.006

Botvinik-Nezer, R., Holzmeister, F., Camerer, C.F., Dreber, A., Huber, J., Johannesson, M., Kirchler, M., Iwanir, R., Mumford, J.A., Adcock, R.A., Avesani, P., Baczkowski, B.M., Bajracharya, A., Bakst, L., Ball, S., Barilari, M., Bault, N., Beaton, D., Beitner, J., Benoit, R.G., Berkers, R.M.W.J., Bhanji, J.P., Biswal, B.B., Bobadilla-Suarez, S., Bortolini, T., Bottenhorn, K.L., Bowring, A., Braem, S., Brooks, H.R., Brudner, E.G., Calderon, C.B., Camilleri, J.A., Castrellon, J.J., Cecchetti, L., Cieslik, E.C., Cole, Z.J., Collignon, O., Cox, R.W., Cunningham, W.A., Czoschke, S., Dadi, K., Davis, C.P., Luca, A.D., Delgado, M.R., Demetriou, L., Dennison, J.B., Di, X., Dickie, E.W., Dobryakova, E., Donnat, C.L., Dukart, J., Duncan, N.W., Durnez, J., Eed, A., Eickhoff, S.B., Erhart, A., Fontanesi, L., Fricke, G.M., Fu, S., Galván, A., Gau, R., Genon, S., Glatard, T., Glerean, E., Goeman, J.J., Golowin, S.A.E., González-García, C., Gorgolewski, K.J., Grady, C.L., Green, M.A., Guassi Moreira, J.F., Guest, O., Hakimi, S., Hamilton, J.P., Hancock, R., Handjaras, G., Harry, B.B., Hawco, C., Herholz, P., Herman, G., Heunis, S., Hoffstaedter, F., Hogeveen, J., Holmes, S., Hu, C.-P., Huettel, S.A., Hughes, M.E., Iacovella, V., Iordan, A.D., Isager, P.M., Isik, A.I., Jahn, A., Johnson, M.R., Johnstone, T., Joseph, M.J.E., Juliano, A.C., Kable, J.W., Kassinopoulos, M., Koba, C., Kong, X.-Z., Koscik, T.R., Kucukboyaci, N.E., Kuhl, B.A., Kupek, S., Laird, A.R., Lamm, C., Langner, R., Lauharatanahirun, N., Lee, H., Lee, S., Leemans, A., Leo, A., Lesage, E., Li, F., Li, M.Y.C., Lim, P.C., Lintz, E.N., Liphardt, S.W., Losecaat Vermeer, A.B., Love, B.C., Mack, M.L., Malpica, N., Marins, T., Maumet, C., McDonald, K., McGuire, J.T., Melero, H., Méndez Leal, A.S., Meyer, B., Meyer, K.N., Mihai, G., Mitsis, G.D., Moll, J., Nielson, D.M., Nilsonne, G., Notter, M.P., Olivetti, E., Onicas, A.I., Papale, P., Patil, K.R., Peelle, J.E., Pérez, A., Pischedda, D., Poline, J.-B., Prystauka, Y., Ray, S., Reuter-Lorenz, P.A., Reynolds, R.C., Ricciardi, E., Rieck, J.R., Rodriguez-Thompson, A.M., Romyn, A., Salo, T., Samanez-Larkin, G.R., Sanz-Morales, E., Schlichting, M.L., Schultz, D.H., Shen, Q., Sheridan, M.A., Silvers, J.A., Skagerlund, K., Smith, A., Smith, D.V., Sokol-Hessner, P., Steinkamp, S.R., Tashjian, S.M., Thirion, B., Thorp, J.N., Tinghög, G., Tisdall, L., Tompson, S.H., Toro-Serey, C., Torre Tresols, J.J., Tozzi, L., Truong, V., Turella, L., van ’t Veer, A.E., Verguts, T., Vettel, J.M., Vijayarajah, S., Vo, K., Wall, M.B., Weeda, W.D., Weis, S., White, D.J., Wisniewski, D., Xifra-Porxas, A., Yearling, E.A., Yoon, S., Yuan, R., Yuen, K.S.L., Zhang, L., Zhang, X., Zosky, J.E., Nichols, T.E., Poldrack, R.A., Schonberg, T., 2020. Variability in the analysis of a single neuroimaging dataset by many teams. Nature 582, 84–88. https://doi.org/10.1038/s41586-020-2314-9

Breakspear, M., 2017. Dynamic models of large-scale brain activity. Nat. Neurosci. 20, 340–352. https://doi.org/10.1038/nn.4497

Bruckmann, S., Hauk, D., Roessner, V., Resch, F., Freitag, C.M., Kammer, T., Ziemann, U., Rothenberger, A., Weisbrod, M., Bender, S., 2012. Cortical inhibition in attention deficit hyperactivity disorder: new insights from the electroencephalographic response to transcranial magnetic stimulation. Brain J. Neurol. 135, 2215–2230. https://doi.org/10.1093/brain/aws071

Casarotto, S., Fecchio, M., Rosanova, M., Varone, G., D’Ambrosio, S., Sarasso, S., Pigorini, A., Russo, S., Comanducci, A., Ilmoniemi, R.J., Massimini, M., 2021. The rt-TEP tool: real-time visualization of TMS-Evoked Potential to maximize cortical activation and minimize artifacts. https://doi.org/10.1101/2021.09.15.460488

Casula, E.P., Bertoldo, A., Tarantino, V., Maiella, M., Koch, G., Rothwell, J.C., Toffolo, G.M., Bisiacchi, P.S., 2017. TMS-evoked long-lasting artefacts: A new adaptive algorithm for EEG signal correction. Clin. Neurophysiol. Off. J. Int. Fed. Clin. Neurophysiol. 128, 1563–1574. https://doi.org/10.1016/j.clinph.2017.06.003

Chung, S.W., Lewis, B.P., Rogasch, N.C., Saeki, T., Thomson, R.H., Hoy, K.E., Bailey, N.W., Fitzgerald, P.B., 2017. Demonstration of short-term plasticity in the dorsolateral prefrontal cortex with theta burst stimulation: A TMS-EEG study. Clin. Neurophysiol. Off. J. Int. Fed. Clin. Neurophysiol. 128, 1117–1126. https://doi.org/10.1016/j.clinph.2017.04.005

Cline, C.C., Lucas, M.V., Sun, Y., Menezes, M., Etkin, A., 2021. Advanced Artifact Removal for Automated TMS-EEG Data Processing, in: 2021 10th International IEEE/EMBS Conference on Neural Engineering (NER). Presented at the 2021 10th International IEEE/EMBS Conference on Neural Engineering (NER), pp. 1039–1042. https://doi.org/10.1109/NER49283.2021.9441147

Cona, F., Zavaglia, M., Massimini, M., Rosanova, M., Ursino, M., 2011. A neural mass model of interconnected regions simulates rhythm propagation observed via TMS-EEG. NeuroImage 57, 1045–1058. https://doi.org/10.1016/j.neuroimage.2011.05.007

Conde, V., Tomasevic, L., Akopian, I., Stanek, K., Saturnino, G.B., Thielscher, A., Bergmann, T.O., Siebner, H.R., 2019. The non-transcranial TMS-evoked potential is an inherent source of ambiguity in TMS-EEG studies. NeuroImage 185, 300–312. https://doi.org/10.1016/j.neuroimage.2018.10.052

Cracco, R.Q., Amassian, V.E., Maccabee, P.J., Cracco, J.B., 1989. Comparison of human transcallosal responses evoked by magnetic coil and electrical stimulation. Electroencephalogr. Clin. Neurophysiol. 74, 417–424. https://doi.org/10.1016/0168-5597(89)90030-0

Daskalakis, Z.J., Farzan, F., Barr, M.S., Maller, J.J., Chen, R., Fitzgerald, P.B., 2008. Long-interval cortical inhibition from the dorsolateral prefrontal cortex: a TMS-EEG study. Neuropsychopharmacol. Off. Publ. Am. Coll. Neuropsychopharmacol. 33, 2860–2869. https://doi.org/10.1038/npp.2008.22

de Cheveigné, A., Nelken, I., 2019. Filters: When, Why, and How (Not) to Use Them. Neuron 102, 280–293. https://doi.org/10.1016/j.neuron.2019.02.039

Delorme, A., Makeig, S., 2004. EEGLAB: an open source toolbox for analysis of single-trial EEG dynamics including independent component analysis. J. Neurosci. Methods 134, 9–21. https://doi.org/10.1016/j.jneumeth.2003.10.009

Farzan, F., Vernet, M., Shafi, M.M.D., Rotenberg, A., Daskalakis, Z.J., Pascual-Leone, A., 2016. Characterizing and Modulating Brain Circuitry through Transcranial Magnetic Stimulation Combined with Electroencephalography. Front. Neural Circuits 10, 73. https://doi.org/10.3389/fncir.2016.00073

Fernandez, L., Biabani, M., Do, M., Opie, G.M., Hill, A.T., Barham, M.P., Teo, W.-P., Byrne, L.K., Rogasch, N.C., Enticott, P.G., 2021. Assessing cerebellar-cortical connectivity using concurrent TMS-EEG: a feasibility study. J. Neurophysiol. 125, 1768–1787. https://doi.org/10.1152/jn.00617.2020

Ferrarelli, F., Sarasso, S., Guller, Y., Riedner, B.A., Peterson, M.J., Bellesi, M., Massimini, M., Postle, B.R., Tononi, G., 2012. Reduced natural oscillatory frequency of frontal thalamo- cortical circuits in schizophrenia. Arch. Gen. Psychiatry 69, 766–774. https://doi.org/10.1001/archgenpsychiatry.2012.147

Fitzgibbon, S.P., DeLosAngeles, D., Lewis, T.W., Powers, D.M.W., Grummett, T.S., Whitham, E.M., Ward, L.M., Willoughby, J.O., Pope, K.J., 2016. Automatic determination of EMG-contaminated components and validation of independent component analysis using EEG during pharmacologic paralysis. Clin. Neurophysiol. Off. J. Int. Fed. Clin. Neurophysiol. 127, 1781–1793. https://doi.org/10.1016/j.clinph.2015.12.009

Freche, D., Naim-Feil, J., Peled, A., Levit-Binnun, N., Moses, E., 2018. A quantitative physical model of the TMS-induced discharge artifacts in EEG. PLoS Comput. Biol. 14, e1006177. https://doi.org/10.1371/journal.pcbi.1006177

Gordon, P.C., Desideri, D., Belardinelli, P., Zrenner, C., Ziemann, U., 2018. Comparison of cortical EEG responses to realistic sham versus real TMS of human motor cortex. Brain Stimulat. 11, 1322–1330. https://doi.org/10.1016/j.brs.2018.08.003

Gordon, P.C., Jovellar, D.B., Song, Y., Zrenner, C., Belardinelli, P., Siebner, H.R., Ziemann, U., 2021. Recording brain responses to TMS of primary motor cortex by EEG - utility of an optimized sham procedure. NeuroImage 245, 118708. https://doi.org/10.1016/j.neuroimage.2021.118708

Gorgolewski, K.J., Auer, T., Calhoun, V.D., Craddock, R.C., Das, S., Duff, E.P., Flandin, G., Ghosh, S.S., Glatard, T., Halchenko, Y.O., Handwerker, D.A., Hanke, M., Keator, D., Li, X., Michael, Z., Maumet, C., Nichols, B.N., Nichols, T.E., Pellman, J., Poline, J.-B., Rokem, A., Schaefer, G., Sochat, V., Triplett, W., Turner, J.A., Varoquaux, G., Poldrack, R.A., 2016. The brain imaging data structure, a format for organizing and describing outputs of neuroimaging experiments. Sci. Data 3, 160044. https://doi.org/10.1038/sdata.2016.44

Habibollahi Saatlou, F., Rogasch, N.C., McNair, N.A., Biabani, M., Pillen, S.D., Marshall, T.R., Bergmann, T.O., 2018. MAGIC: An open-source MATLAB toolbox for external control of transcranial magnetic stimulation devices. Brain Stimulat. 11, 1189–1191. https://doi.org/10.1016/j.brs.2018.05.015

Hamidi, M., Slagter, H.A., Tononi, G., Postle, B.R., 2010. Brain responses evoked by high-frequency repetitive transcranial magnetic stimulation: an event-related potential study. Brain Stimulat. 3, 2–14. https://doi.org/10.1016/j.brs.2009.04.001

Hassan, U., Pillen, S., Zrenner, C., Bergmann, T.O., 2021. The Brain Electrophysiological recording & STimulation (BEST) toolbox. Brain Stimulat. 15, 109–115. https://doi.org/10.1016/j.brs.2021.11.017

Hernandez-Pavon, J.C., Metsomaa, J., Mutanen, T., Stenroos, M., Mäki, H., Ilmoniemi, R.J., Sarvas, J., 2012. Uncovering neural independent components from highly artifactual TMS-evoked EEG data. J. Neurosci. Methods 209, 144–157. https://doi.org/10.1016/j.jneumeth.2012.05.029

Herring, J.D., Thut, G., Jensen, O., Bergmann, T.O., 2015. Attention Modulates TMS-Locked Alpha Oscillations in the Visual Cortex. J. Neurosci. Off. J. Soc. Neurosci. 35, 14435–14447. https://doi.org/10.1523/JNEUROSCI.1833-15.2015

Hyvarinen, A., 1999. Fast and robust fixed-point algorithms for independent component analysis. IEEE Trans. Neural Netw. 10, 626–634. https://doi.org/10.1109/72.761722

Ilmoniemi, R.J., Hernandez-Pavon, J.C., Makela, N.N., Metsomaa, J., Mutanen, T.P., Stenroos, M., Sarvas, J., 2015. Dealing with artifacts in TMS-evoked EEG. Annu. Int. Conf. IEEE Eng. Med. Biol. Soc. IEEE Eng. Med. Biol. Soc. Annu. Int. Conf. 2015, 230–233. https://doi.org/10.1109/EMBC.2015.7318342

Ilmoniemi, R.J., Kicić, D., 2010. Methodology for combined TMS and EEG. Brain Topogr. 22, 233–248. https://doi.org/10.1007/s10548-009-0123-4

Ilmoniemi, R.J., Virtanen, J., Ruohonen, J., Karhu, J., Aronen, H.J., Näätänen, R., Katila, T., 1997. Neuronal responses to magnetic stimulation reveal cortical reactivity and connectivity. Neuroreport 8, 3537–3540. https://doi.org/10.1097/00001756-199711100-00024

Izumi, S., Takase, M., Arita, M., Masakado, Y., Kimura, A., Chino, N., 1997. Transcranial magnetic stimulation-induced changes in EEG and responses recorded from the scalp of healthy humans. Electroencephalogr. Clin. Neurophysiol. 103, 319–322. https://doi.org/10.1016/s0013-4694(97)00007-2

Julkunen, P., Pääkkönen, A., Hukkanen, T., Könönen, M., Tiihonen, P., Vanhatalo, S., Karhu, J., 2008. Efficient reduction of stimulus artefact in TMS-EEG by epithelial short-circuiting by mini-punctures. Clin. Neurophysiol. Off. J. Int. Fed. Clin. Neurophysiol. 119, 475–481. https://doi.org/10.1016/j.clinph.2007.09.139

Korhonen, R.J., Hernandez-Pavon, J.C., Metsomaa, J., Mäki, H., Ilmoniemi, R.J., Sarvas, J., 2011. Removal of large muscle artifacts from transcranial magnetic stimulation-evoked EEG by independent component analysis. Med. Biol. Eng. Comput. 49, 397–407. https://doi.org/10.1007/s11517-011-0748-9

Mäki, H., Ilmoniemi, R.J., 2011. Projecting out muscle artifacts from TMS-evoked EEG. NeuroImage 54, 2706–2710. https://doi.org/10.1016/j.neuroimage.2010.11.041

Mancuso, M., Sveva, V., Cruciani, A., Brown, K., Ibáñez, J., Rawji, V., Casula, E., Premoli, I., D’Ambrosio, S., Rothwell, J., Rocchi, L., 2021. Transcranial Evoked Potentials Can Be Reliably Recorded with Active Electrodes. Brain Sci. 11, 145. https://doi.org/10.3390/brainsci11020145

Markiewicz, C.J., Gorgolewski, K.J., Feingold, F., Blair, R., Halchenko, Y.O., Miller, E., Hardcastle, N., Wexler, J., Esteban, O., Goncavles, M., Jwa, A., Poldrack, R., 2021. The OpenNeuro resource for sharing of neuroscience data. eLife 10, e71774. https://doi.org/10.7554/eLife.71774

Massimini, M., Ferrarelli, F., Huber, R., Esser, S.K., Singh, H., Tononi, G., 2005. Breakdown of cortical effective connectivity during sleep. Science 309, 2228–2232. https://doi.org/10.1126/science.1117256

Metsomaa, J., Sarvas, J., Ilmoniemi, R.J., 2014. Multi-trial evoked EEG and independent component analysis. J. Neurosci. Methods 228, 15–26. https://doi.org/10.1016/j.jneumeth.2014.02.019

Mutanen, T., Mäki, H., Ilmoniemi, R.J., 2013. The effect of stimulus parameters on TMS-EEG muscle artifacts. Brain Stimulat. 6, 371–376. https://doi.org/10.1016/j.brs.2012.07.005

Mutanen, T.P., Biabani, M., Sarvas, J., Ilmoniemi, R.J., Rogasch, N.C., 2020. Source-based artifact-rejection techniques available in TESA, an open-source TMS-EEG toolbox. Brain Stimulat. 13, 1349–1351. https://doi.org/10.1016/j.brs.2020.06.079

Mutanen, T.P., Kukkonen, M., Nieminen, J.O., Stenroos, M., Sarvas, J., Ilmoniemi, R.J., 2016. Recovering TMS-evoked EEG responses masked by muscle artifacts. NeuroImage 139, 157–166. https://doi.org/10.1016/j.neuroimage.2016.05.028

Mutanen, T.P., Metsomaa, J., Liljander, S., Ilmoniemi, R.J., 2018. Automatic and robust noise suppression in EEG and MEG: The SOUND algorithm. NeuroImage 166, 135–151. https://doi.org/10.1016/j.neuroimage.2017.10.021

Nikouline, V., Ruohonen, J., Ilmoniemi, R.J., 1999. The role of the coil click in TMS assessed with simultaneous EEG. Clin. Neurophysiol. Off. J. Int. Fed. Clin. Neurophysiol. 110, 1325–1328. https://doi.org/10.1016/s1388-2457(99)00070-x

Nikulin, V.V., Nolte, G., Curio, G., 2011. A novel method for reliable and fast extraction of neuronal EEG/MEG oscillations on the basis of spatio-spectral decomposition. NeuroImage 55, 1528–1535. https://doi.org/10.1016/j.neuroimage.2011.01.057

Oostenveld, R., Fries, P., Maris, E., Schoffelen, J.-M., 2011. FieldTrip: Open source software for advanced analysis of MEG, EEG, and invasive electrophysiological data. Comput. Intell. Neurosci. 2011, 156869. https://doi.org/10.1155/2011/156869

Ozdemir, R.A., Tadayon, E., Boucher, P., Sun, H., Momi, D., Ganglberger, W., Westover, M.B., Pascual-Leone, A., Santarnecchi, E., Shafi, M.M., 2021. Cortical responses to noninvasive perturbations enable individual brain fingerprinting. Brain Stimulat. 14, 391–403. https://doi.org/10.1016/j.brs.2021.02.005

Paus, T., Sipila, P.K., Strafella, A.P., 2001. Synchronization of neuronal activity in the human primary motor cortex by transcranial magnetic stimulation: an EEG study. J. Neurophysiol. 86, 1983–1990. https://doi.org/10.1152/jn.2001.86.4.1983

Poldrack, R.A., Gorgolewski, K.J., 2014. Making big data open: data sharing in neuroimaging. Nat. Neurosci. 17, 1510–1517. https://doi.org/10.1038/nn.3818

Rocchi, L., Di Santo, A., Brown, K., Ibáñez, J., Casula, E., Rawji, V., Di Lazzaro, V., Koch, G., Rothwell, J., 2021. Disentangling EEG responses to TMS due to cortical and peripheral activations. Brain Stimulat. 14, 4–18. https://doi.org/10.1016/j.brs.2020.10.011

Rogasch, N.C., Daskalakis, Z.J., Fitzgerald, P.B., 2015. Cortical inhibition of distinct mechanisms in the dorsolateral prefrontal cortex is related to working memory performance: A TMS–EEG study. Cortex 64, 68–77. https://doi.org/10.1016/j.cortex.2014.10.003

Rogasch, N.C., Fitzgerald, P.B., 2013. Assessing cortical network properties using TMS-EEG. Hum. Brain Mapp. 34, 1652–1669. https://doi.org/10.1002/hbm.22016

Rogasch, N.C., Sullivan, C., Thomson, R.H., Rose, N.S., Bailey, N.W., Fitzgerald, P.B., Farzan, F., Hernandez-Pavon, J.C., 2017. Analysing concurrent transcranial magnetic stimulation and electroencephalographic data: A review and introduction to the open-source TESA software. NeuroImage 147, 934–951. https://doi.org/10.1016/j.neuroimage.2016.10.031

Rogasch, N.C., Thomson, R.H., Daskalakis, Z.J., Fitzgerald, P.B., 2013. Short-latency artifacts associated with concurrent TMS-EEG. Brain Stimulat. 6, 868–876. https://doi.org/10.1016/j.brs.2013.04.004

Rogasch, N.C., Thomson, R.H., Farzan, F., Fitzgibbon, B.M., Bailey, N.W., Hernandez-Pavon, J.C., Daskalakis, Z.J., Fitzgerald, P.B., 2014. Removing artefacts from TMS-EEG recordings using independent component analysis: importance for assessing prefrontal and motor cortex network properties. NeuroImage 101, 425–439. https://doi.org/10.1016/j.neuroimage.2014.07.037

Rogasch, N.C., Zipser, C., Darmani, G., Mutanen, T.P., Biabani, M., Zrenner, C., Desideri, D., Belardinelli, P., Müller-Dahlhaus, F., Ziemann, U., 2020. The effects of NMDA receptor blockade on TMS-evoked EEG potentials from prefrontal and parietal cortex. Sci. Rep. 10, 3168. https://doi.org/10.1038/s41598-020-59911-6

Rosanova, M., Casali, A., Bellina, V., Resta, F., Mariotti, M., Massimini, M., 2009. Natural frequencies of human corticothalamic circuits. J. Neurosci. Off. J. Soc. Neurosci. 29, 7679–7685. https://doi.org/10.1523/JNEUROSCI.0445-09.2009

Ruddy, K.L., Woolley, D.G., Mantini, D., Balsters, J.H., Enz, N., Wenderoth, N., 2018. Improving the quality of combined EEG-TMS neural recordings: Introducing the coil spacer. J. Neurosci. Methods 294, 34–39. https://doi.org/10.1016/j.jneumeth.2017.11.001

Russo, S., Sarasso, S., Puglisi, G.E., Palù, D.D., Pigorini, A., Casarotto, S., D’Ambrosio, S., Astolfi, A., Massimini, M., Rosanova, M., Fecchio, M., 2021. TAAC - TMS Adaptable Auditory Control: a universal tool to mask TMS click. https://doi.org/10.1101/2021.09.08.459439

Salo, K.S.-T., Mutanen, T.P., Vaalto, S.M.I., Ilmoniemi, R.J., 2020. EEG Artifact Removal in TMS Studies of Cortical Speech Areas. Brain Topogr. 33, 1–9. https://doi.org/10.1007/s10548-019-00724-w

Sanz-Leon, P., Robinson, P.A., Knock, S.A., Drysdale, P.M., Abeysuriya, R.G., Fung, F.K., Rennie, C.J., Zhao, X., 2018. NFTsim: Theory and Simulation of Multiscale Neural Field Dynamics. PLoS Comput. Biol. 14, e1006387. https://doi.org/10.1371/journal.pcbi.1006387

Schwarz, C.G., Kremers, W.K., Therneau, T.M., Sharp, R.R., Gunter, J.L., Vemuri, P., Arani, A., Spychalla, A.J., Kantarci, K., Knopman, D.S., Petersen, R.C., Jack, C.R., 2019. Identification of Anonymous MRI Research Participants with Face-Recognition Software. N. Engl. J. Med. 381, 1684–1686. https://doi.org/10.1056/NEJMc1908881

Sekiguchi, H., Takeuchi, S., Kadota, H., Kohno, Y., Nakajima, Y., 2011. TMS-induced artifacts on EEG can be reduced by rearrangement of the electrode’s lead wire before recording. Clin. Neurophysiol. Off. J. Int. Fed. Clin. Neurophysiol. 122, 984–990. https://doi.org/10.1016/j.clinph.2010.09.004

Siebner, H.R., Conde, V., Tomasevic, L., Thielscher, A., Bergmann, T.O., 2019. Distilling the essence of TMS-evoked EEG potentials (TEPs): A call for securing mechanistic specificity and experimental rigor. Brain Stimulat. 12, 1051–1054. https://doi.org/10.1016/j.brs.2019.03.076

Silberzahn, R., Uhlmann, E.L., Martin, D.P., Anselmi, P., Aust, F., Awtrey, E., Bahník, Š., Bai, F., Bannard, C., Bonnier, E., Carlsson, R., Cheung, F., Christensen, G., Clay, R., Craig, M.A., Dalla Rosa, A., Dam, L., Evans, M.H., Flores Cervantes, I., Fong, N., Gamez- Djokic, M., Glenz, A., Gordon-McKeon, S., Heaton, T.J., Hederos, K., Heene, M., Hofelich Mohr, A.J., Högden, F., Hui, K., Johannesson, M., Kalodimos, J., Kaszubowski, E., Kennedy, D.M., Lei, R., Lindsay, T.A., Liverani, S., Madan, C.R., Molden, D., Molleman, E., Morey, R.D., Mulder, L.B., Nijstad, B.R., Pope, N.G., Pope, B., Prenoveau, J.M., Rink, F., Robusto, E., Roderique, H., Sandberg, A., Schlüter, E., Schönbrodt, F.D., Sherman, M.F., Sommer, S.A., Sotak, K., Spain, S., Spörlein, C., Stafford, T., Stefanutti, L., Tauber, S., Ullrich, J., Vianello, M., Wagenmakers, E.-J., Witkowiak, M., Yoon, S., Nosek, B.A., 2018. Many Analysts, One Data Set: Making Transparent How Variations in Analytic Choices Affect Results. Adv. Methods Pract. Psychol. Sci. 1, 337–356. https://doi.org/10.1177/2515245917747646

Stokes, M.G., Chambers, C.D., Gould, I.C., Henderson, T.R., Janko, N.E., Allen, N.B., Mattingley, J.B., 2005. Simple metric for scaling motor threshold based on scalp-cortex distance: application to studies using transcranial magnetic stimulation. J. Neurophysiol. 94, 4520–4527. https://doi.org/10.1152/jn.00067.2005

Tadel, F., Baillet, S., Mosher, J.C., Pantazis, D., Leahy, R.M., 2011. Brainstorm: a user-friendly application for MEG/EEG analysis. Comput. Intell. Neurosci. 2011, 879716. https://doi.org/10.1155/2011/879716

ter Braack, E.M., de Jonge, B., van Putten, M.J.A.M., 2013. Reduction of TMS induced artifacts in EEG using principal component analysis. IEEE Trans. Neural Syst. Rehabil. Eng. Publ. IEEE Eng. Med. Biol. Soc. 21, 376–382. https://doi.org/10.1109/TNSRE.2012.2228674

Tremblay, S., Rogasch, N.C., Premoli, I., Blumberger, D.M., Casarotto, S., Chen, R., Di Lazzaro, V., Farzan, F., Ferrarelli, F., Fitzgerald, P.B., Hui, J., Ilmoniemi, R.J., Kimiskidis, V.K., Kugiumtzis, D., Lioumis, P., Pascual-Leone, A., Pellicciari, M.C., Rajji, T., Thut, G., Zomorrodi, R., Ziemann, U., Daskalakis, Z.J., 2019. Clinical utility and prospective of TMS-EEG. Clin. Neurophysiol. Off. J. Int. Fed. Clin. Neurophysiol. 130, 802–844. https://doi.org/10.1016/j.clinph.2019.01.001

Van Der Werf, Y.D., Paus, T., 2006. The neural response to transcranial magnetic stimulation of the human motor cortex. I. Intracortical and cortico-cortical contributions. Exp. Brain Res. 175, 231–245. https://doi.org/10.1007/s00221-006-0551-2

Veniero, D., Bortoletto, M., Miniussi, C., 2009. TMS-EEG co-registration: on TMS-induced artifact. Clin. Neurophysiol. Off. J. Int. Fed. Clin. Neurophysiol. 120, 1392–1399. https://doi.org/10.1016/j.clinph.2009.04.023

Virtanen, J., Ruohonen, J., Näätänen, R., Ilmoniemi, R.J., 1999. Instrumentation for the measurement of electric brain responses to transcranial magnetic stimulation. Med. Biol. Eng. Comput. 37, 322–326. https://doi.org/10.1007/BF02513307

White, T., Blok, E., Calhoun, V.D., 2020. Data sharing and privacy issues in neuroimaging research: Opportunities, obstacles, challenges, and monsters under the bed. Hum. Brain Mapp. https://doi.org/10.1002/hbm.25120

Winkler, I., Debener, S., Müller, K.-R., Tangermann, M., 2015. On the influence of high-pass filtering on ICA-based artifact reduction in EEG-ERP. Annu. Int. Conf. IEEE Eng. Med. Biol. Soc. IEEE Eng. Med. Biol. Soc. Annu. Int. Conf. 2015, 4101–4105. https://doi.org/10.1109/EMBC.2015.7319296

Wu, W., Keller, C.J., Rogasch, N.C., Longwell, P., Shpigel, E., Rolle, C.E., Etkin, A., 2018. ARTIST: A fully automated artifact rejection algorithm for single-pulse TMS-EEG data. Hum. Brain Mapp. 39, 1607–1625. https://doi.org/10.1002/hbm.23938

Zanon, M., Busan, P., Monti, F., Pizzolato, G., Battaglini, P.P., 2010. Cortical connections between dorsal and ventral visual streams in humans: Evidence by TMS/EEG co-registration. Brain Topogr. 22, 307–317. https://doi.org/10.1007/s10548-009-0103-8

